# Predicting fungal secondary metabolite activity from biosynthetic gene cluster data using machine learning

**DOI:** 10.1101/2023.09.12.557468

**Authors:** Olivia Riedling, Allison S. Walker, Antonis Rokas

## Abstract

Fungal secondary metabolites (SMs) play a significant role in the diversity of ecological communities, niches, and lifestyles in the fungal kingdom. Many fungal SMs have medically and industrially important properties including antifungal, antibacterial, and antitumor activity, and a single metabolite can display multiple types of bioactivities. The genes necessary for fungal SM biosynthesis are typically found in a single genomic region forming biosynthetic gene clusters (BGCs). However, whether fungal SM bioactivity can be predicted from specific attributes of genes in BGCs remains an open question. We adapted previously used machine learning models for predicting SM bioactivity from bacterial BGC data to fungal BGC data. We trained our models to predict antibacterial, antifungal, and cytotoxic/antitumor bioactivity on two datasets: 1) fungal BGCs (dataset comprised of 314 BGCs), and 2) fungal (314 BGCs) and bacterial BGCs (1,003 BGCs); the second dataset was our control since a previous study using just the bacterial BGC data yielded prediction accuracies as high as 80%. We found that the models trained only on fungal BGCs had balanced accuracies between 51-68%, whereas training on bacterial and fungal BGCs yielded balanced accuracies between 61-74%. The lower accuracy of the predictions from fungal data likely stems from the small number of BGCs and SMs with known bioactivity; this lack of data currently limits the application of machine learning approaches in studying fungal secondary metabolism. However, our data also suggest that machine learning approaches trained on bacterial and fungal data can predict SM bioactivity with good accuracy. With more than 15,000 characterized fungal SMs, millions of putative BGCs present in fungal genomes, and increased demand for novel drugs, efforts that systematically link fungal SM bioactivity to BGCs are urgently needed.

## Introduction

Fungi have captivated the scientific community for centuries due to their diverse ecological roles and remarkable ability to produce an array of bioactive secondary metabolites (SMs). SMs are small biologically active compounds that aid in adapting to different environments but are not required for normal function or survival^1^. Many clades across the fungal kingdom produce SMs with a large amount of them belonging to the *Pezizomycotina* subphylum of filamentous fungi.

These SMs have diverse bioactivities, including as antifungals^2^, UV protectants^3^, antibacterials^4–6^, iron sequesterers^7^, antifeedants^8^, immunosuppressants^9^, and toxins^10^. In addition to their ecological importance, SMs are also of considerable interest to medicine, industry, and the bioeconomy, and are used in a wide range of applications. Many SMs such as penicillin^11^, an antibiotic, and lovastatin^12^, a cholesterol-lowering antifungal, are iconic drugs. Other SMs have been utilized in the food science, agricultural, and cosmetic industries for bioactivities such as antibiotics, pigments, and antifeedants.

The genes involved in fungal SM production are typically located in a single genomic region and are known as biosynthetic gene clusters (BGCs)^1,13^. It is estimated that fungal genomes may have up to 30-70 BGCs^14^, sometimes even more; these BGCs contain many different types of genes, such as self-resistance genes whose protein products confer protection against the produced SMs, transport genes, transcriptional regulatory genes, core or backbone genes, whose protein products synthesize the backbone of the SM, and backbone modifying genes^1^. There is a variety of core genes, such as polyketide synthases (PKS), terpene synthases, and non-ribosomal peptide synthases (NRPS)^14^. The genes within these BGCs often vary considerably in terms of gene presence, absence, and orientation within and between species^13,15^ contributing to fungal ecological and chemical diversity^1^.

Identification of novel fungal SMs is a labor-intensive process consisting of isolation, purification, and structural elucidation of novel compounds^16^. Novel fungal are typically subjected to bioassays for the detection of specific biological activities (e.g., antifungal or antibacterial)^17^. Unfortunately, there is currently no systematic effort for gathering fungal SM bioactivity data, limiting their potential utility in large scale analyses. More recently, discovery of novel SMs in diverse organisms, including bacteria, plants, and fungi, has been accelerated by advances in genomic sequencing, genetic engineering, and bioinformatics. Advances in omics technologies have enabled discovery of novel BGC-SM pairs through genomic manipulations and the utilization of heterologous expression to isolate SMs from unculturable species^18–20^, and activation or increase in SM production through promoter engineering^21^.

The increasing number of characterized BGC-SM pairs from diverse organisms has led to the creation of large repositories such as the Minimum Information about a Biosynthetic Gene Cluster (MIBiG) database^22^, which houses standardized annotations and metadata on BGC-SM pairs, increasing the efficiency of natural product discovery and facilitating additional analyses. In parallel, novel computational approaches, such as artificial intelligence or machine learning, are being employed to predict BGCs^23^, like ClusterFinder^24^, or predict both BGCs and SM bioactivities, like DeepBGC^25^. The ability to predict fungal SM bioactivity will allow further study of SMs across the fungal kingdom and generate candidate SMs for drug discovery. It has been observed that fungal SM chemical properties are “more drug-like” compared to their bacterial counterparts by US-FDA guidelines^26^, and the deep well of fungal derived SMs has yet to be fully taken advantage of. However, whether fungal SM bioactivity can be predicted from specific features of genes in BGCs remains an open question.

In this work, we adapted machine learning models by Walker and Clardy^27^ that predict bacterial SM bioactivity from bacterial BGC data with accuracies as high as 80%^27^ to test whether we could predict fungal SM bioactivity from fungal BGC data. We trained the models to predict antibacterial, antifungal, and cytotoxic/antitumor SM activity from BGC data, using two training datasets: 1) fungal BGC data, and 2) fungal and bacterial BGC data (as a control, since predictions based on training using bacterial BGC data previously yielded relatively high accuracies^27^). The models trained on the fungal dataset (comprised of 314 BGCs) had balanced accuracies between 51-68%, which were lower than the balanced accuracies between 61-74% of models trained on the fungal and bacterial dataset (comprised of 314 fungal and 1,003 bacterial BGCs). The lower balanced accuracies in the models trained on fungal data compared to the models by Walker and Clardy^27^ are likely due to limited data available on characterized fungal BGCs and SMs. In contrast, models trained on both fungal and bacterial data can predict SM bioactivity with relatively good accuracies. We conclude that the small number of fungal BGC-SM pairs with known bioactivities currently limits the use of artificial intelligence approaches for predicting SM bioactivity. Given this limitation, we recommend that both bacterial and fungal data are used in machine learning training data and call for efforts that systematically catalog fungal SM bioactivity (including lack of bioactivity) and identify new BGC-SM pairs.

## Methods

### Obtaining fungal and bacterial BGC-SM pairs with known bioactivities

The fungal BGC GenBank files of 392 fungal BGCs were downloaded from the MIBiG database, version 3.0^22^. We performed a literature search of bioactivity and growth assays to identify validated bioactivities of the SM products from all 392 fungal BGCs in MIBiG. Data on the 1,152 bacterial BGCs and their SM product bioactivities were retrieved from the study by Walker and Clardy^27^. SM bioactivities were categorized into “antibacterial”, “antifungal”, “cytotoxic”, “antitumor”, and “unknown” for when data were not available; we also identified SM bioactivities that were not included in our predictions (e.g., antifeedant) because of their small numbers. Bioactivity classifications were converted into binary matrices, such that “1” indicated validated presence of activity and “0” indicated absence of activity or lack of documentation. These data are available in Table S1.

### Feature selection and construction of training datasets

Our training data included the BGC number (corresponding to the accession number of the BGC in the MIBiG database), product name, the SM product bioactivity, core gene present in the BGC that biosynthesizes the backbone of the SM product, and the species and genus classification. BGCs whose bioactivities are unknown were not included in model training. Two training datasets were used: the first contained only fungal BGCs and the bioactivities of their corresponding SMs (392 BGCs in total; the SMs of 123 BGCs had antibacterial bioactivity, of 115 had antifungal bioactivity, of 96 had cytotoxic bioactivity, and of 123 had antitumor bioactivity) and the second contained both fungal (392 BGCs) and bacterial BGCs (1,544 BGCs in total; the SMs of 627 BGCs had antibacterial bioactivity, of 312 had antifungal bioactivity, of 328 had cytotoxic bioactivity, and of 260 had antitumor bioactivity) and the bioactivities of their SMs. From the 392 BGCs in the fungal dataset, 78 were removed because their bioactivities were unknown, resulting in 314 BGCs used in training the models; similarly, from the 1,544 BGCs in the fungal and bacterial dataset. 227 were removed because their bioactivities were unknown, leaving 1,317 BGCs (314 fungal and 1,003 bacterial) used in training the models.

To obtain the features for model training, each GenBank file for the BGCs was run through antiSMASH, version 5^7^. For each BGC, we extracted from the GenBank output files generated by antiSMASH: 1) Pfam protein family domains represented in the BGC, 2) the core gene(s) (PKS, NRPS, etc.,), 3) cluster-defining CDS features (gene features that antiSMASH uses to define the BGC class or classes (e.g., NRPS, PKS, etc.)), and 4) annotations of secondary metabolite clusters of orthologous groups of proteins (smCOGs, annotations for accessory genes in BGCs based on sequence similarity to genes in other characterized BGCs). Extractions were preformed using Python scripts modified from Walker and Clardy^27^. To identify genes in the BGCs similar to antibiotic resistance genes, which have the potential to be predictive of antibacterial and antifungal bioactivity, the GenBank files generated by antiSMASH, one for each BGC, were converted into fasta files and subsequently run through Resistance Gene Identifier (RGI), (version 5)^29^. The RGI annotations were extracted similarly to the antiSMASH output with a Python script, retaining only the genes that occurred five or more times in the dataset. All the features collected were converted into binary matrices and used in model training.

### Machine Learning Models

To gain insights into fungal SM bioactivity we trained machine learning models to predict three types of bioactivity: antibacterial, antifungal, and cytotoxic/antitumor. Following Walker and Clardy^27^, we used three algorithms to make the three binary classifications: 1) Stochastic Gradient Decent Classifier (SGDClassifier) module for the logistic regression (LR), 2) random forest (RF) with extra randomized decision trees, and 3) the Support Vector Clustering (SVC) module for support vector machine (SVM). All three classifiers were used independently to predict each type of bioactivity since a SM can have multiple bioactivity types. Each classifier was imported from the scikit-learn Python library^30^, and parameters were determined by completing GridSearch from scikit-learn and then a 10-fold cross-validation to evaluate the accuracy of each classifier. The parameter values used in the GridSearch for the LR models were maximum iterations of 100; log loss, elasticnet penalties; alpha values of .5, .3, .2, .1, .01, .001, .0001, .00001, .000001; and l1 ratios of .5, .2, .05, .1, .01, .001, and .0001. Additionally, the tolerance was set to “none”. The parameter values used in the GridSearch for the SVM models were c-values of 100, 10, 1, .5, .1, and .01 and gamma values of .01, .1, 1, and 10. The parameter values for the RF models were maximum features on auto; Gini criterion; bootstraps true; maximum depth of 10, 20, 50, 100, 1000, and none; and number of estimators were 1, 5, 10, 15, 25, 50, and 100. The best parameter values for each classifier were chosen based on the highest average accuracy.

To evaluate the models, we used the balanced accuracy metric from the scikit-learn Python library. We used balanced accuracy to address the imbalance of the bioactivity types in the training data since the bioactivities are rare and therefore have a larger portion of the 0’s compared to 1’s (e.g., there are 116 BGCs with antifungal bioactivity (1’s) compared to 198 without (0’s) in our fungal BGC training data). Each of the classifiers was assessed in 10-fold cross validation, and the balanced accuracy of each classifier was compared to classifiers trained on randomized features that represent the inability to distinguish between classes (0 or 1). Each classifier for each classification was assessed using a one-way ANOVA to determine if there was a significant difference between using the actual training data vs. the randomized data. All the ANOVAs were performed through the Python library SciPy^31^ and assessed using alpha levels of 0.0001, 0.001, 0.01, and 0.05. Classifiers in each classification problem were also evaluated with receiver operator characteristic (ROC) curves and precision-recall (PR) curves. ROC curves plot the true positive rate against the false positive rate. Classifiers with the largest area under the curve (AUC) values have better true-to-false positive ratios, with AUC values greater than 0.5 indicating better than random ability to correctly predict the presence/absence of a given bioactivity. The precision-recall (PR) curves plot the recall (i.e., true positives over the sum of true positives and false negatives) against the precision (i.e., true positives over the sum of the true positives and the false positives); they are considered more accurate for classifiers trained on imbalanced datasets, such as ours.

### Assembling the fungal phylogeny

To analyze the distribution of the characterized SM bioactivities across the fungal kingdom we displayed the different types of bioactivity known for different species on the branch tips of a phylogeny using the iToL (Interactive Tree of Life) tool, version 6^32^. The phylogeny was modified with no tip labels and no branch lengths from a previous phylogenomic analysis of 290 genes from 1,644 fungal species^33^. If a species in the dataset was not present in the phylogeny, we chose a close relative in the same genus; if the genus was also absent, the data for that species were not displayed on the phylogeny.

## Data Availability

All input files and scripts required to reproduce the results in this study will become available upon acceptance of the manuscript for publication.

## Results and Discussion

### Classifiers trained on only fungal data have lower balanced accuracies

To predict fungal SM bioactivity from BGC data, we trained three machine learning classifiers on only fungal BGCs and features (245 features from 314 fungal BGCs with known bioactivities out of 392 total BGCs in the dataset). The distribution of bioactivities in the training data were 39% antibacterial, 37% antifungal, and 56% cytotoxic/antitumor. The ANOVA analyses comparing the balanced accuracies of the classifiers trained on training data vs. randomized features were significant in all cases apart from the LR for antibacterial predictions (**Figure 1A**). The balanced accuracies for antifungal (SVM: 66%, LR: 64%, RF: 68%) were the highest followed by cytotoxic/antitumor (SVM: 59%, LR: 58%, RF: 61%) and antibacterial (SVM: 55%, LR: 51%, RF: 58%) classifications (**Figure 1A**).

**Figure 1:**
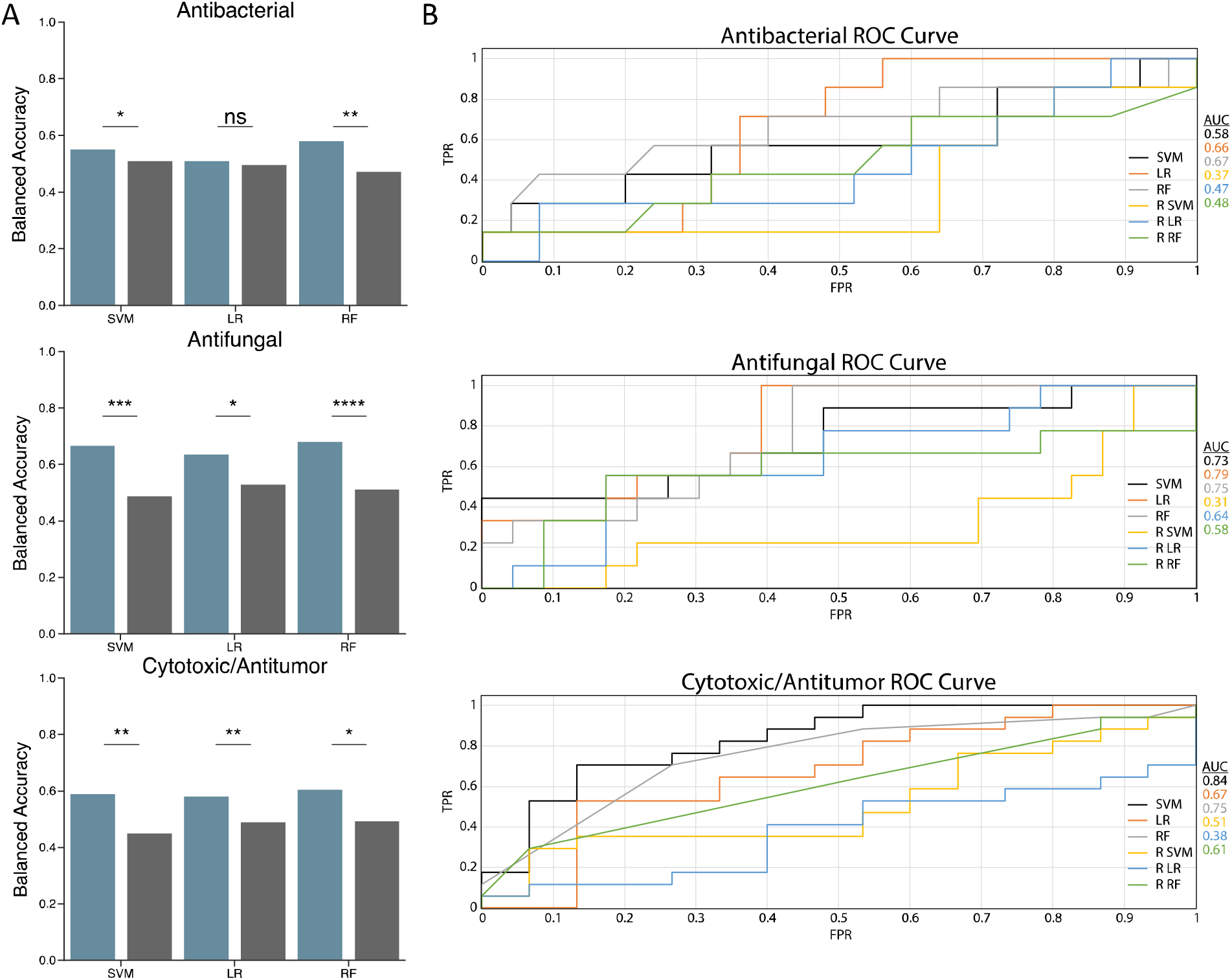
Machine learning models trained only on fungal BGC data exhibited low balanced accuracies. A) Balanced accuracy of classifiers. X-axis shows support vector machine (SVM), logistic regression (LR), and random forest (RF) classifiers trained on actual data (blue), and classifiers trained on randomized features (grey). Y-axis shows the balanced accuracy of classifiers. Stars indicate significance of one-way ANOVA at 0.05(*), 0.01(**), 0.001(***), and 0.0001(****). B) Receiver Operator Characteristic (ROC) curves for all classifiers. X-axis shows the false positive rate (FPR), and the y- axis shows the true positive rate (TPR). Lines of different colors correspond to the three classifiers trained on actual data (SVM, LR, and RF) and to the three classifiers trained on randomized data (R SVM, R LR, R RF). Area under the curve (AUC) is shown to the right for each classifier.

To identify potential explanations for the low balanced accuracies observed in our classifications, we examined the PR (better at evaluating datasets with moderate to large class imbalances), and ROC curves (better at evaluating datasets where classes are equal and balanced). Additionally, we analyzed the rates of true positives (TPR), false positives (FPR), true negatives (TNR), and false negatives (FNR) in each of our classifications. We found that the AUCs for the PR curves were consistently lower than the ROC curves for the antibacterial and antifungal classifications (**Figure 1B**) (**Figure S1**). This indicated a low number of false positives (explaining why the ROC curves, which plot the FPR against the TPR, but do not account for false negatives, were higher) and a larger number of false negatives (explaining why the PR curves, which plot recall or the true positives/true positives + false negatives against the precision or the true positives/true positives + false positives were lower). The larger number of false negatives in the antibacterial and antifungal classifications, which is also shown in the confusion matrices from the cross-validation and the average FPR and FNR values (**Figures S3-4**) (**Table S2**), suggest that the classifiers are training on the negative class (i.e., on the 0’s) instead of the positive class (1’s) (**Figure S1**). Even though the class of SMs with antibacterial bioactivity is larger than the antifungal class in the training data, the antibacterial classifiers had lower balanced accuracies and higher FNRs compared to the antifungal classifiers. This result could be due to a larger number of shared features between BGCs in the antifungal bioactivity class that increased its prediction accuracy compared to the antibacterial bioactivity class (**Figure S6**). The cytotoxic/antitumor classification had relatively similar ROC and PR AUCs and had high FPRs and TPRs and low FNRs and TNRs (**Table S2**). This result suggests that the classifiers are training on the positive class (1’s) but overclassifying 0’s as 1’s, potentially due the larger number of 1’s for the cytotoxic/antitumor bioactivity in the fungal dataset (177 SMs with cytotoxic/antitumor bioactivity out of 314 total) (**Figure S7**). In summary, the relatively low accuracies for all three classifications are likely due to our small, imbalanced training dataset.

### Classifiers trained on fungal and bacterial data have higher balanced accuracies

The classifiers trained on fungal data displayed relatively low accuracies compared to classifiers trained on bacterial data from a previous study^27^. Thus, we next trained classifiers on both fungal and bacterial data from a combined dataset. If the bacterial and fungal BGCs have shared features that correlate with the bioactivity of their SMs in the same way, the combined dataset should have increased balanced accuracies. However, if there are not enough shared features or they do not correlate to bioactivity, training on the combined dataset will result in a decrease in balanced accuracies.

There were 1,152 bacterial BGCs in the training data, but 149 BGCs had unknown bioactivities, so the final bacterial data included 1,003 bacterial BGCs. Combining the 1,003 bacterial BGCs with the 314 fungal BGCs resulting in a training dataset comprised of 1,317 BGCs and 984 features. The distribution of bioactivities in the training data were antibacterials: 47%, antifungals: 24%, and 35% cytotoxic/antitumor. The ANOVA analyses comparing the balanced accuracies of classifiers trained on the training data vs. on randomized data were all significant (p<0.0001) (**Figure 2A**). The balanced accuracies were between ∼61% to∼74%, with the antibacterial classification having the highest balanced accuracies (SVM: 74%, LR: 72%, RF: 74%) followed by cytotoxic/antitumor (SVM: 70%, LR: 68%, RF: 73%), and antifungal (SVM: 64%, LR: 61%, RF: 66%) classifications.

**Figure 2:**
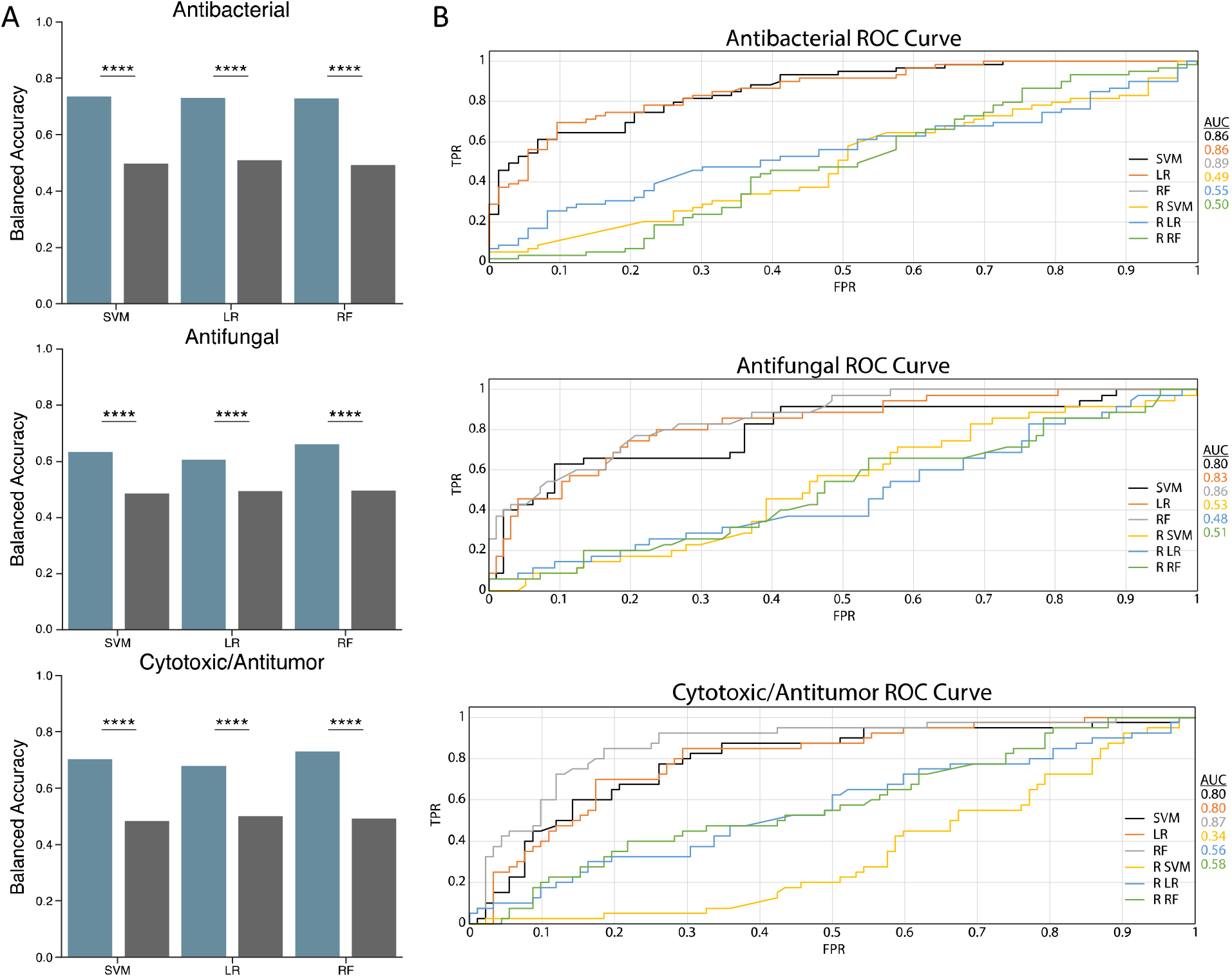
Classifiers trained on both fungal and bacterial BGC data exhibit higher balanced accuracies. A) Balanced accuracy of classifiers. X-axis shows support vector machine (SVM), logistic regression (LR), and random forest (RF) classifiers trained on actual data (blue), and classifiers trained on randomized features (grey). Y-axis shows the balanced accuracy of classifiers. Stars indicate significance of one-way ANOVA at 0.05(*), 0.01(**), 0.001(***), and 0.0001(****). B) Receiver Operator Characteristic (ROC) curves for all classifiers. X-axis shows the false positive rate (FPR), and the y- axis shows the true positive rate (TPR). Lines of different colors correspond to the three classifiers trained on actual data (SVM, LR, and RF) and to the three classifiers trained on randomized data (R SVM, R LR, R RF). Area under the curve (AUC) is shown to the right for each classifier.

Our balanced accuracies are slightly lower compared to the original study by Walker and Clardy^27^, where the models were trained on just the bacterial BGC data. The observed differences are potentially due to the differing types of features, such as PFAM domains and cluster-defining features, between fungal and bacterial BGCs, and the imbalanced distributions of bioactivity types in the training data. Walker and Clardy also used sequence similarity networks which added an additional 825 features for the classifiers trained on bacterial data in their study. However, the removal of these features did not significantly impact accuracies. Compared to the balanced accuracies based on only fungal BGC data, the balanced accuracies based on bacterial and fungal data increased, suggesting that the features in the bacterial and fungal BGCs are similar enough to increase prediction accuracy.

We examined the ROC and PR curves to identify reasons for the reduced accuracy relative to Walker and Clardy. The ROC curves for all classifications have high AUCs with the antibacterial classification (SVM: 0.86, LR: 0.86, RF: 0.89) being the highest, followed by the antifungal (SVM: 0.80, LR: 0.83, RF: 0.86) and cytotoxic/antitumor (SVM: 0.80, LR: 0.80, RF: 0.87) classifications (**Figure 2B**). The antibacterial classification had ROC AUC (SVM: 0.86, LR: 0.86, RF: 0.89) values that were similar to the PR AUC (SVM:0.85, LR: 0.85, RF:0.89) values, and the antifungal classification had ROC AUC (SVM: 0.80, LR: 0.83, RF: 0.86) values that were higher than the PR AUC (SVM:0.64, LR:0.66, RF:0.73) values. These results show that the classifiers in the antibacterial classification have better performance than the antifungal classifiers when trained on both bacterial and fungal data vs. when trained on only fungal data.

This is likely due to the much larger number of SMs with antibacterial bioactivity in the fungal and bacterial training dataset. In contrast, the cytotoxic/antitumor classification had ROC AUCs (SVM: 0.80, LR: 0.80, RF: 0.87) that were substantially higher than PR AUCs (SVM:0.56, LR:0.56, RF:0.72) (**Figure S2**). In general, lower PR AUC values indicate poor performance on the positive class. To explain the lower PR AUC values compared to the ROC AUC values in the cytotoxic/antitumor classification, we analyzed the FPR, TPR, FNR, and TNR. The average FNR for the cytotoxic/antitumor classification was relatively high compared to the FPR (**Table S3**) and is likely the reason for the reduction in the PR AUC values as seen in the cross-validation confusion matrices (**Figure S9**). The high FNR in the cytotoxic/antitumor classification could be due to the lack of shared features between fungal and bacterial BGCs.

### Lower balanced accuracies in stem from the lack of fungal BGC data

An additional explanation for the lower accuracies of our models trained on fungal and bacterial data relative to the accuracies observed by Walker and Clardy with training on only bacterial data could be that there is substantial diversity in features that contribute to each bioactivity between bacterial and fungal BGCs. For example, there are numerous antibacterial agents that do not share the same targets; some have bacteriostatic or bactericidal properties but have the same mode of inhibition in preventing protein synthesis^34^, while others inhibit peptidoglycan biosynthesis to form pores in bacterial membranes. Additionally, some have broad spectrum or narrow bioactivity against different microbes increasing their specificity. The small size of our dataset, especially of fungal BGCs, likely does not sufficiently capture the ways in which different BGC features can affect bioactivity for different mechanisms of actions, especially for those that have multiple bioactivity types. Consistent with this explanation, feature importance analysis showed that the most informative features in the antibacterial and cytotoxic/antitumor bioactivity predictions were very different between the classifiers trained on fungal data and classifiers trained on fungal and bacterial data (**Figures S10-11**).

In contrast, the most informative features in the antifungal bioactivity predictions had a large amount of overlap between the two training datasets. A few of the features that appeared in both datasets were crotonyl-CoA reductase, AMP-binding enzyme, multiple polyketide synthase domains, and fungal specific transcription factor domain (**Figure 3**). This overlap of features that exhibit antifungal bioactivities between fungal and bacterial BGCs may stem from many antifungals targeting the cell membrane or pathways involved in its assembly (e.g., ergosterol biosynthesis)^35^.

**Figure 3:**
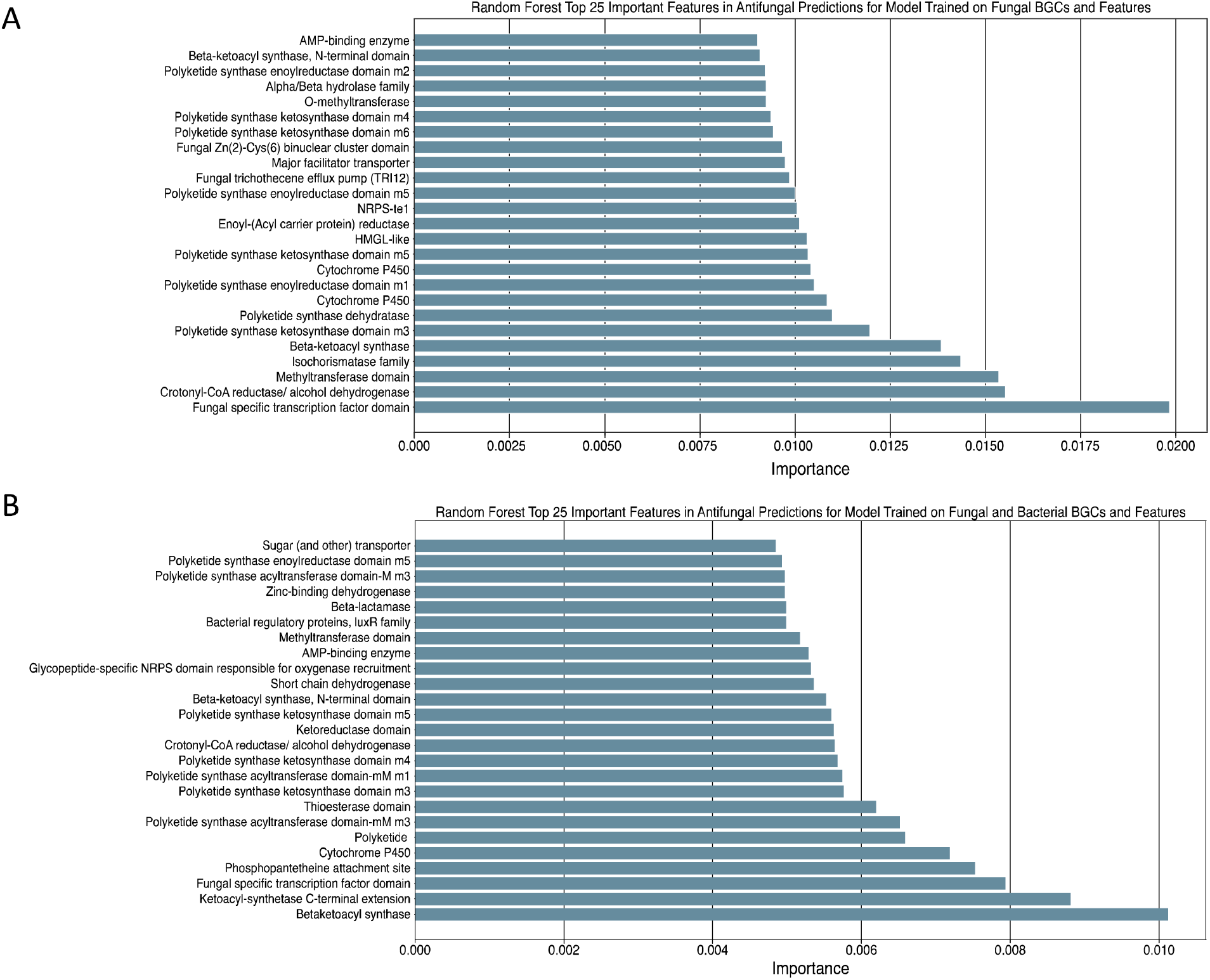
The 25 most important features in antifungal predictions are similar between models trained on just fungal BGC data vs. models trained on fungal and bacterial BGC data. A) Top 25 important features in antifungal predictions for the models trained on fungal data. X-axis shows importance and Y-axis shows the features. B) Top 25 important features in antifungal predictions for the models trained on fungal and bacterial data. X-axis shows importance and Y-axis shows the features.

Nevertheless, there were also features that were specific to the fungal dataset. For example, in models trained on only fungal data the PFAM identifiers HMGL-like and isochorismatase family were present in ∼5% of the BGCs with antifungal bioactivity and in 0% of the BGCs without.

The HGML-like domain encompasses a family of aldolases and a region of pyruvate carboxylases that all contain phosphate binding sites, and the isochorismatase family is a family of hydrolase enzymes. Both enzymes are involved in conversion steps in metabolic pathways and isochorismatase family enzymes have been involved in the degradation of creatinine in *Pseudomonas putida* and *Anthrobacter* sp.^36^. Additionally, the polyketide synthase ketosynthase domain in module 3 (PKSI-KS m3) of BGCs was present in ∼78% of the BGCs with antifungal bioactivity vs. ∼54% of BGCs without, and the beta-ketoacyl synthase, N-terminal domain was present in 79% of BGCs with antifungal bioactivity vs. ∼56% of BGCs without in models trained on only fungal data.

### Narrow taxonomic distribution of SM bioactivities across the fungal kingdom

Another potential explanation for the low accuracies of the classifiers trained on fungal BGC data may be that the bioactivities of already characterized fungal SMs stem from only a subset of fungal clades. To analyze the distribution of characterized SM bioactivity types across the fungal kingdom as well as determine if there are any phylogenetic patterns in the bioactivities we studied, we mapped the bioactivity types to a phylogeny of 1,644 species that span the diversity of the fungal kingdom^33^. The 314 fungal BGCs used in our training data stemmed from: 78 species that produced BGCs with SM products that had antibacterial bioactivity (53 were in the phylogeny, 5 were not in the phylogeny, and we used another representative from the same genus for 20 species), 81 with antifungal bioactivity (47 in the phylogeny, 7 not in the phylogeny, we used another representative from the same genus for 27 species), 62 with cytotoxic bioactivity (35 in the phylogeny, 5 not in the phylogeny, we used another representative from the genus for 22 species), and 79 with antitumor bioactivity (45 in the phylogeny, 8 not in the phylogeny, we used another representative from same genus for 24 species).

While there were not any notable phylogenetic patterns in the distributions of bioactivity types across fungi, there was a very sporadic distribution of characterized BGC-SM pairs with areas of relatively dense coverage (**Figure 4**). A few genera were well sampled compared to the rest of the phylogeny, including *Aspergillus*, *Penicillium,* and *Fusarium*; in contrast, BGC-SM pairs with characterized bioactivity were rather sparsely distributed across most of the phylogeny. This was the case even in the *Pezizomycotina* subphylum of filamentous fungi, which are well known to contain many BGCs^37^.

**Figure 4:**
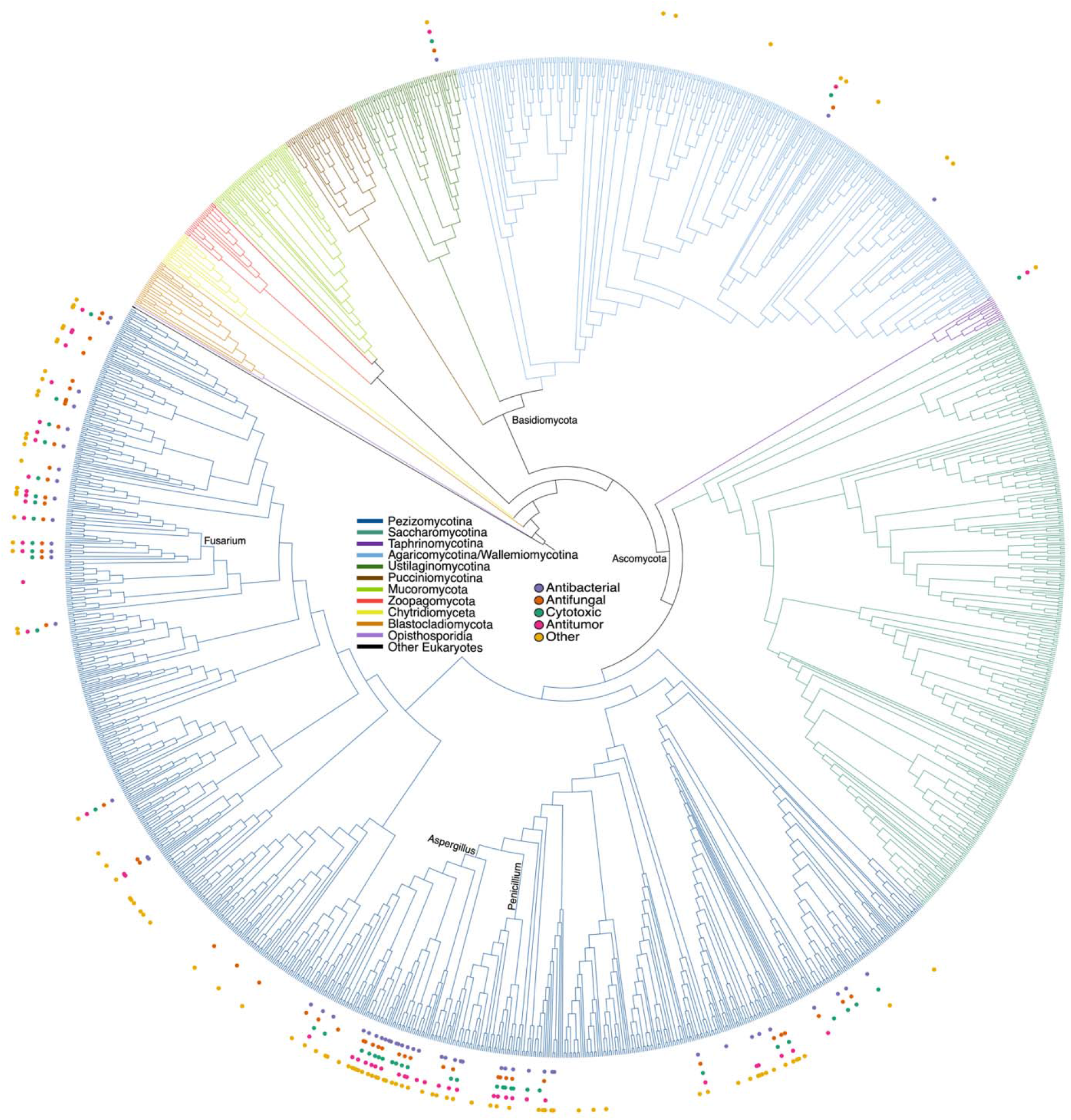
Narrow taxonomic distribution of characterized biosynthetic gene cluster-secondary metabolite pairs across the fungal kingdom. Phylogeny modified from Li et al. 2021^33^. Branch colors indicate different subphyla within the fungal kingdom. Circles indicate the four activity types (antibacterial, antifungal, antitumor, cytotoxic) included in the predictions and other activity types noted in dataset (e.g., antifeedant) as well as the secondary metabolites with unknown bioactivity (others).

We analyzed the distribution of genera in the fungal dataset and observed that while few genera contained many BGC-SM pairs, there were many that had none or only one representative in the dataset (**Figure S12A**). Additionally, of the 392 BGC SM pairs in the original dataset and the 314 fungal BGCs used in model training, 131 and 101, respectively, were derived from the genus *Aspergillus*, indicating a large bias in characterization of fungal BGC-SM pairs. Notably, this taxonomic bias is not unique to fungal BGCs-SM pairs. We observed a very similar distribution in the bacterial data, where over 400 of the bacterial BGCs-SM pairs are from the genus *Streptomyces* (**Figure S13**).

We next used antiSMASH to determine if the lower number of BGC-SM pairs in certain fungal species or genera was due to smaller number of BGCs in their genomes. We found that the number of predicted BGCs across species did not correlate with the number of BGC-SM pairs, suggesting that the lack of representation of certain taxa in the dataset is likely due to lack of studies that link these BGCs to their corresponding SMs and their bioactivities (**Figure S12B**).

Given the bias in characterization is clearly depicted in the number of BGC-SM pairs derived from *Aspergillus* and *Streptomyces*, we next examined whether the BGC features from these two genera were representative of the BGC features in the rest of fungi and bacteria, respectively. We analyzed the proportion of features present in *Aspergillus* and *Streptomyces* BGCs and compared it to the proportion of features present in non-*Aspergillus* and non-*Streptomyces* derived BGCs. In the fungal dataset, we observed that the proportions of features between *Aspergillus* derived BGCs and non-*Aspergillus* derived BGCs were similar (**Figure 5**). This suggests that the features present in *Aspergillus* BGCs are likely representative of the diversity of features in the entire fungal dataset. We observed a similar result in the dataset with bacterial and fungal BGCs with notably more features absent in *Streptomyces* BGCs than in the non-*Streptomyces* BGCs (**Figure S14**). These results suggest that the taxonomic bias in BGC-SM pair characterization does not significantly bias the feature types present in the data. Furthermore, in the fungal data specifically, these results suggest that the low accuracy in using machine learning to predict fungal SM bioactivity is due the small number of characterized fungal BGC-SM pairs rather than due to the lack of diversity of BGCs from different species in the training data. In other words, both *Aspergillus* and *Streptomyces* genera are potentially good models to use in machine learning due the current numbers of characterized BGC-SM pairs and the types of features present.

**Figure 5:**
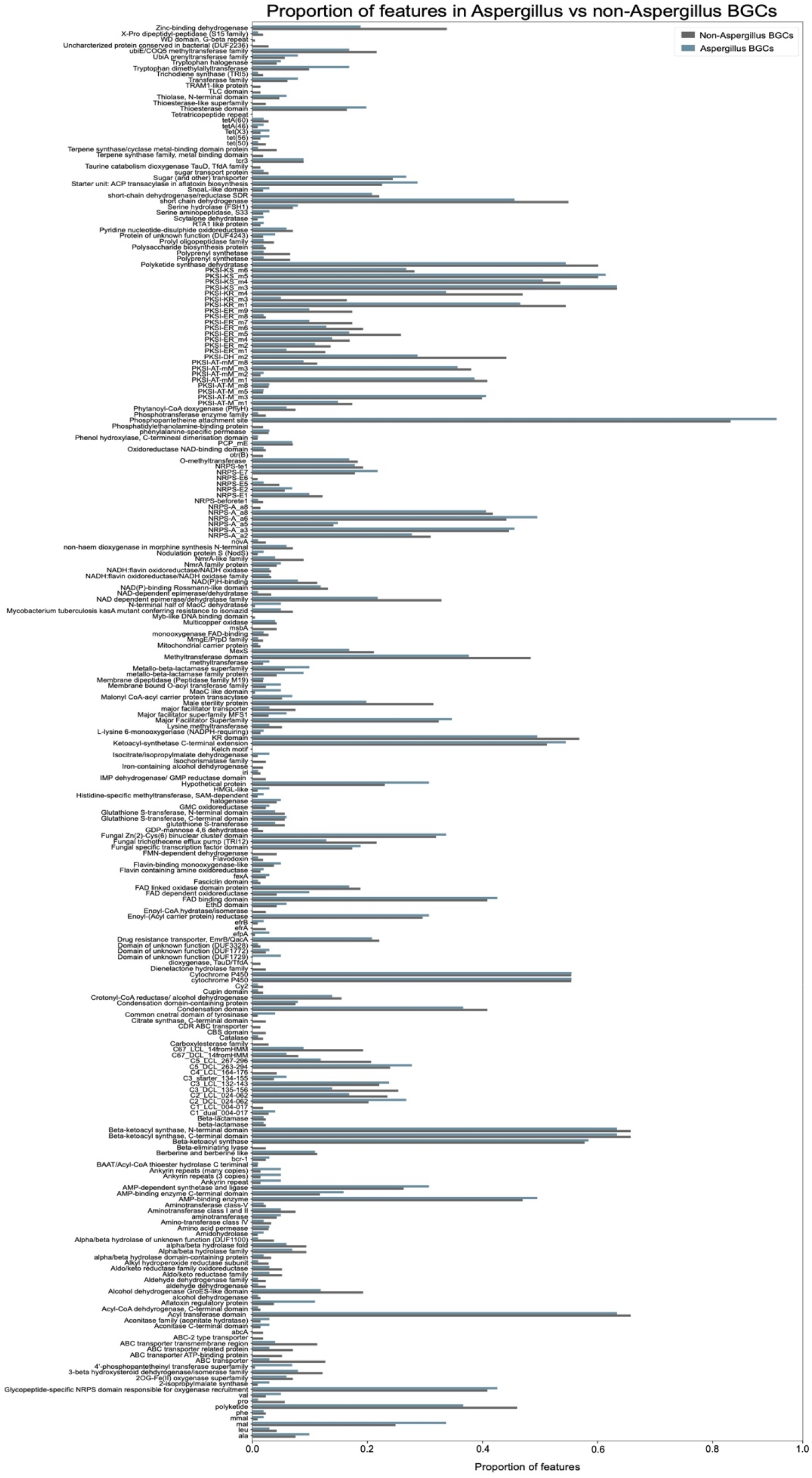
Features in *Aspergillus* BGCs are representative of the diversity of features in the entire fungal dataset. X-axis shows the proportion of feature presence in *Aspergillus* (blue) vs. *non-Aspergillus* (grey) BGCs, Y-axis shows all the features in the fungal dataset.

### The need for systematic effort in characterizing BGC-SM pairs

Currently, there have been more than 15,000 fungal SMs characterized and millions of putative BGCs identified in fungal genomes^38^. Due to the increased demand for novel drugs, efforts that systematically link fungal SM bioactivity to BGCs are urgently needed. Various methods of correlating SMs to their corresponding BGCs have been undertaken such as targeted genome mining, heterologous expression, metabologenomics with gene cluster family (GCF) networking and correlation-based scoring^39^, and feature-based correlation methods utilizing genome-metabolome ontologies in bacterial species^40^. While these methods are continuing to improve, SM discovery is still far ahead of the characterization of the BGCs responsible for their biosynthesis. Although there are various databases available for characterized SMs, such as the Dictionary of Natural Products and Medicinal Fungi Secondary metabolites And Therapeutics (MeFSAT)^41^, there remains potentially millions of uncharacterized BGC-SM pairs. Publishing results of SM bioactivity assays in repositories, including the negative results, would be another systematic effort that could enable larger analyses and greater accessibility to these data. Only a small portion of SMs has been linked to BGCs and an even small portion of these BGC-SM pairs has characterized bioactivities (**Figure S14**). Considering more than 15,000 fungal SMs known to date and new methodologies for linking SMs and BGCs^18^, potential opportunities and benefits of filling this large gap of knowledge are considerable.

Machine learning methods that predict fungal SM bioactivity from BGC data show promise but currently are unable to perform with high accuracy due to the lack of fungal BGC-SM pairs with bioactivity data. Here, we obtained much lower balanced accuracies for models trained on only fungal BGCs (51-68%), likely because only 314 fungal BGCs were used to train our models.

Walker and Clardy^27^ showed that using a dataset with 1,003 bacterial BGCs achieved accuracies as high as ∼80%, so we hypothesize that ∼1,000 fungal BGC-SM pairs with known bioactivities will be a necessary to substantially increase accuracy. Incorporating additional features related to the chemical structures of the SMs could increase the specificity between bioactivity classes and increase the balanced accuracies. A previous study examined the scaffold diversity of fungal SMs^42^, suggesting that complex SM structures could be broken down into different features that capture scaffold content and structural diversity. There are numerous methods of converting chemical structures into machine readable formats like simplified molecular input line entry system representation (SMILES) and molecular fingerprinting, which could enable their direct application into machine learning and advance the understanding of chemical diversity^43^.

## Conclusions

We used machine learning models trained on only fungal BGC data and both fungal and bacterial BGC data to predict SM bioactivity. Due to the lack of available data on fungal BGC-SM pairs, the model trained on only fungal BGC data exhibited relatively low balanced accuracies. The model trained on both fungal and bacterial BGCs shows promise due to the larger amount of BGCs in the dataset, however, further model optimization for predicting fungal SM bioactivity will require a larger dataset of fungal BGC-SM pairs. Additionally, breaking down the large, generalized bioactivity types (antibacterial, antifungal, cytotoxic/antitumor) into more specific classes incorporating the target or mode of action in addition to chemical structure features may aid in more specificity and better accuracies in predicting the bioactivity (although even larger numbers of BGC-SM pairs may be required). Ultimately, improving model overall performance and accuracy will require a systemic effort in characterizing BGC-SM pairs and their bioactivities (including negative results) and depositing the data in large, publicly available repositories. While the current accuracies of artificial intelligence approaches are constrained by the lack of sufficient training data, the potential of machine learning applications in fungal secondary metabolism will remain limited.

## Acknowledgements

We thank members of the Rokas lab for helpful discussion and feedback. Research in A.R.’s lab is supported by grants from the National Science Foundation (DEB-2110404), the National Institutes of Health National Institute of Allergy and Infectious Diseases (R01 AI153356), and the Burroughs Wellcome Fund. Research in A.S.W.’s lab is supported by the National Institutes of Health National Institute of General Medicine (R35 GM146987). The content is solely the responsibility of the authors and does not necessarily represent the official views of the National Institutes of Health.

## References

1. Rokas, A., Mead, M. E., Steenwyk, J. L., Raja, H. A. & Oberlies, N. H. Biosynthetic gene clusters and the evolution of fungal chemodiversity. Nat Prod Rep 37, 868–878 (2020).

2. Brill, G. M., Chen, R. H., Rasmussen, R. R., Whittern, D. N. & Mcalpine, J. B. Calbistrins, novel antifungal agents produced by Penicillium restrictum. II. Isolation and elucidation of structure. J. Antibiot. 46, 39–47 (1993).

3. Nguyen, K.-H., Chollet-Krugler, M., Gouault, N. & Tomasi, S. UV-protectant metabolites from lichens and their symbiotic partners. Nat Prod Rep 30, 1490–1508 (2013).

4. Boudart, G. Antibacterial Activity of Sirodesmin PL Phytotoxin: Application to the Selection of Phytotoxin-Deficient Mutants. Appl Environ Microbiol 55, 1555–1559 (1989).

5. Al-Fakih, A. A. & Almaqtri, W. Q. A. Overview on antibacterial metabolites from terrestrial Aspergillus spp. Mycology 10, 191–209 (2019).

6. Geronikaki, A. et al. Antibacterial activity of griseofulvin analogues as an example of drug repurposing. International Journal of Antimicrobial Agents 55, 105884 (2020).

7. Chen, L.-H., Lin, C.-H. & Chung, K.-R. A nonribosomal peptide synthetase mediates siderophore production and virulence in the citrus fungal pathogen Alternaria alternata. Mol Plant Pathol 14, 497–505 (2013).

8. Stuart, A. E., Brooks, C. J., Prescott, R. J. & Blackwell, A. Repellent and antifeedant activity of salicylic acid and related compounds against the biting midge, Culicoides impunctatus (Diptera: Ceratopogonidae). J Med Entomol 37, 222–227 (2000).

9. Yamada, A., Kataoka, T. & Nagai, K. The fungal metabolite gliotoxin: immunosuppressive activity on CTL-mediated cytotoxicity. Immunol Lett 71, 27–32 (2000).

10. Mamur, S., Ünal, F., Yılmaz, S., Erikel, E. & Yüzbaşıoğlu, D. Evaluation of the cytotoxic and genotoxic effects of mycotoxin fusaric acid. Drug Chem Toxicol 43, 149–157 (2020).

11. Ott, J. L. & Neuss, N. Antibiotic activity of pure penicillin N and isopenicillin N. J. Antibiot. 35, 637–638 (1982).

12. Isolation and characterization of lovastatin producing fungi; investigating the antimicrobial and extracellular enzymatic activities. Int. J. Biosci. 10, 12–20 (2017).

13. Rokas, A., Wisecaver, J. H. & Lind, A. L. The birth, evolution and death of metabolic gene clusters in fungi. Nat Rev Microbiol 16, 731–744 (2018).

14. Keller, N. P. Fungal secondary metabolism: regulation, function and drug discovery. Nat Rev Microbiol 17, 167–180 (2019).

15. Steenwyk, J. L. et al. Variation Among Biosynthetic Gene Clusters, Secondary Metabolite Profiles, and Cards of Virulence Across Aspergillus Species. Genetics 216, 481–497 (2020).

16. Atanasov, A. G. et al. Natural products in drug discovery: advances and opportunities. Nature Reviews Drug Discovery 20, 200–216 (2021).

17. Strömstedt, A. A., Felth, J. & Bohlin, L. Bioassays in Natural Product Research - Strategies and Methods in the Search for Anti-inflammatory and Antimicrobial Activity: ANTIMICROBIAL AND ANTI-INFLAMMATORY ASSAYS IN NATURAL PRODUCT RESEARCH. Phytochem. Anal. 25, 13–28 (2014).

18. Greco, C., Keller, N. P. & Rokas, A. Unearthing fungal chemodiversity and prospects for drug discovery. Curr Opin Microbiol 51, 22–29 (2019).

19. Chiang, Y.-M., Lin, T.-S. & Wang, C. C. C. Total Heterologous Biosynthesis of Fungal Natural Products in Aspergillus nidulans. J. Nat. Prod. 85, 2484–2518 (2022).

20. Zhang, H., Boghigian, B. A., Armando, J. & Pfeifer, B. A. Methods and options for the heterologous production of complex natural products. Nat Prod Rep 28, 125–151 (2011).

21. Umemura, M., Kuriiwa, K., Dao, L. V., Okuda, T. & Terai, G. Promoter tools for further development of Aspergillus oryzae as a platform for fungal secondary metabolite production. Fungal Biol Biotechnol 7, 3 (2020).

22. Medema, M. H. et al. Minimum Information about a Biosynthetic Gene cluster. Nature Chemical Biology 11, 625–631 (2015).

23. Aghdam, S. A. & Brown, A. M. V. Deep learning approaches for natural product discovery from plant endophytic microbiomes. Environmental Microbiome 16, 6 (2021).

24. Cimermancic, P. et al. Insights into Secondary Metabolism from a Global Analysis of Prokaryotic Biosynthetic Gene Clusters. Cell 158, 412–421 (2014).

25. Liu, M., Li, Y. & Li, H. Deep Learning to Predict the Biosynthetic Gene Clusters in Bacterial Genomes. Journal of Molecular Biology 434, 167597 (2022).

26. Robey, M. T., Caesar, L. K., Drott, M. T., Keller, N. P. & Kelleher, N. L. An interpreted atlas of biosynthetic gene clusters from 1,000 fungal genomes. Proc Natl Acad Sci U S A 118, e2020230118 (2021).

27. Walker, A. S. & Clardy, J. A Machine Learning Bioinformatics Method to Predict Biological Activity from Biosynthetic Gene Clusters. J. Chem. Inf. Model. 61, 2560–2571 (2021).

28. Blin, K. et al. antiSMASH 5.0: updates to the secondary metabolite genome mining pipeline. Nucleic Acids Research 47, W81–W87 (2019).

29. McArthur, A. G. et al. The comprehensive antibiotic resistance database. Antimicrob Agents Chemother 57, 3348–3357 (2013).

30. Pedregosa, F. et al. Scikit-Learn: Machine Learning in Python. J. Mach. Learn. Res. 12, 2825–2830 (2011).

31. Virtanen, P. et al. SciPy 1.0: fundamental algorithms for scientific computing in Python. Nature Methods 17, 261–272 (2020).

32. Letunic, I. & Bork, P. Interactive Tree Of Life (iTOL) v5: an online tool for phylogenetic tree display and annotation. Nucleic Acids Research 49, W293–W296 (2021).

33. Li, Y. et al. A genome-scale phylogeny of the kingdom Fungi. Current Biology 31, 1653–1665.e5 (2021).

34. Ullah, H. & Ali, S. Classification of AntiLJBacterial Agents and Their Functions. In *Antibacterial Agents* (ed. Kumavath, R. N.) (InTech, 2017). doi:10.5772/intechopen.68695.

35. Andriole, V. T. Current and future antifungal therapy: new targets for antifungal agents. Journal of Antimicrobial Chemotherapy 44, 151–162 (1999).

36. Romão, M. J., et al. Crystal structure analysis, refinement and enzymatic reaction mechanism of N-carbamoylsarcosine amidohydrolase from Arthrobacter sp. at 2·0Åresolution. Journal of Molecular Biology 226, 1111–1130 (1992).

37. Wisecaver, J. H., Slot, J. C. & Rokas, A. The Evolution of Fungal Metabolic Pathways. PLoS Genet 10, e1004816 (2014).

38. Bills, G. F. & Gloer, J. B. Biologically Active Secondary Metabolites from the Fungi. Microbiol Spectr 4, 4.6.01 (2016).

39. Caesar, L. K. et al. Correlative metabologenomics of 110 fungi reveals metabolite–gene cluster pairs. Nature Chemical Biology 19, 846–854 (2023).

40. Louwen, J. J. R., Medema, M. H. & van der Hooft, J. J. J. Enhanced correlation-based linking of biosynthetic gene clusters to their metabolic products through chemical class matching. Microbiome 11, 13 (2023).

41. Vivek-Ananth, R. P., Sahoo, A. K., Kumaravel, K., Mohanraj, K. & Samal, A. MeFSAT: a curated natural product database specific to secondary metabolites of medicinal fungi. RSC Adv 11, 2596–2607 (2021).

42. González-Medina, M. et al. Scaffold Diversity of Fungal Metabolites. Front. Pharmacol. 8, (2017).

43. Raghunathan, S. & Priyakumar, U. D. Molecular representations for machine learning applications in chemistry. Int J of Quantum Chemistry 122, e26870 (2022).

