## Supplementary Figures for "Predicting fungal secondary metabolite activity from biosynthetic gene cluster data using machine learning"

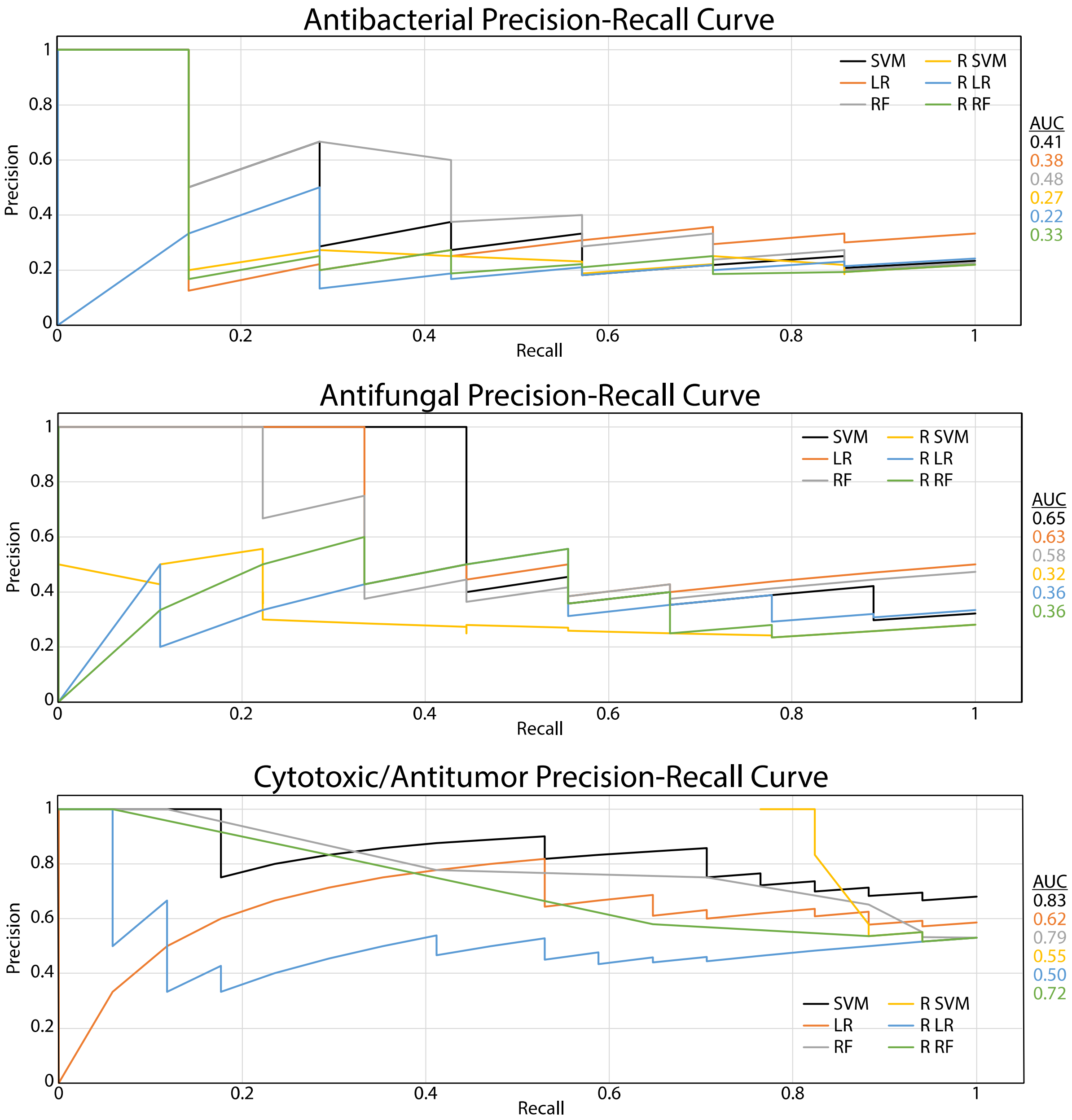


**Supplemental Figure 1**: Precision-Recall Curves for each classification and all three classifiers for models trained on fungal biosynthetic gene clusters (BGCs). X-axis shows the recall (True positives/(True positives + False negatives)) and Y-axis shows the precision (True positives/(True positives + False positives))

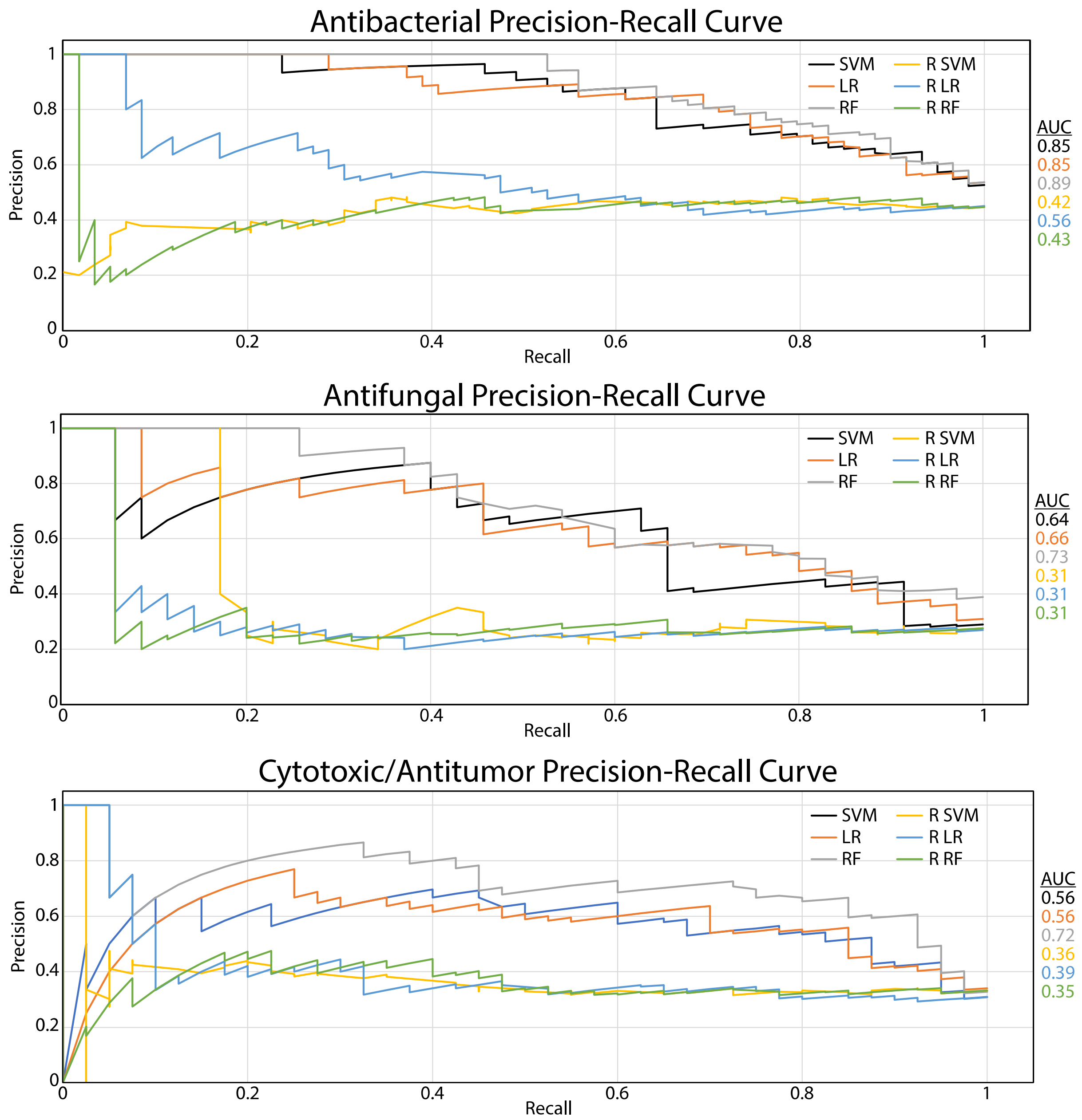


**Supplemental Figure 2**: Precision-Recall Curves for each classification and all three classifiers for models trained on bacterial and fungal biosynthetic gene clusters (BGCs). X-axis shows the recall (True positives/(True positives + False negatives)) and Y-axis shows the precision (True positives/(True positives + False positives)).

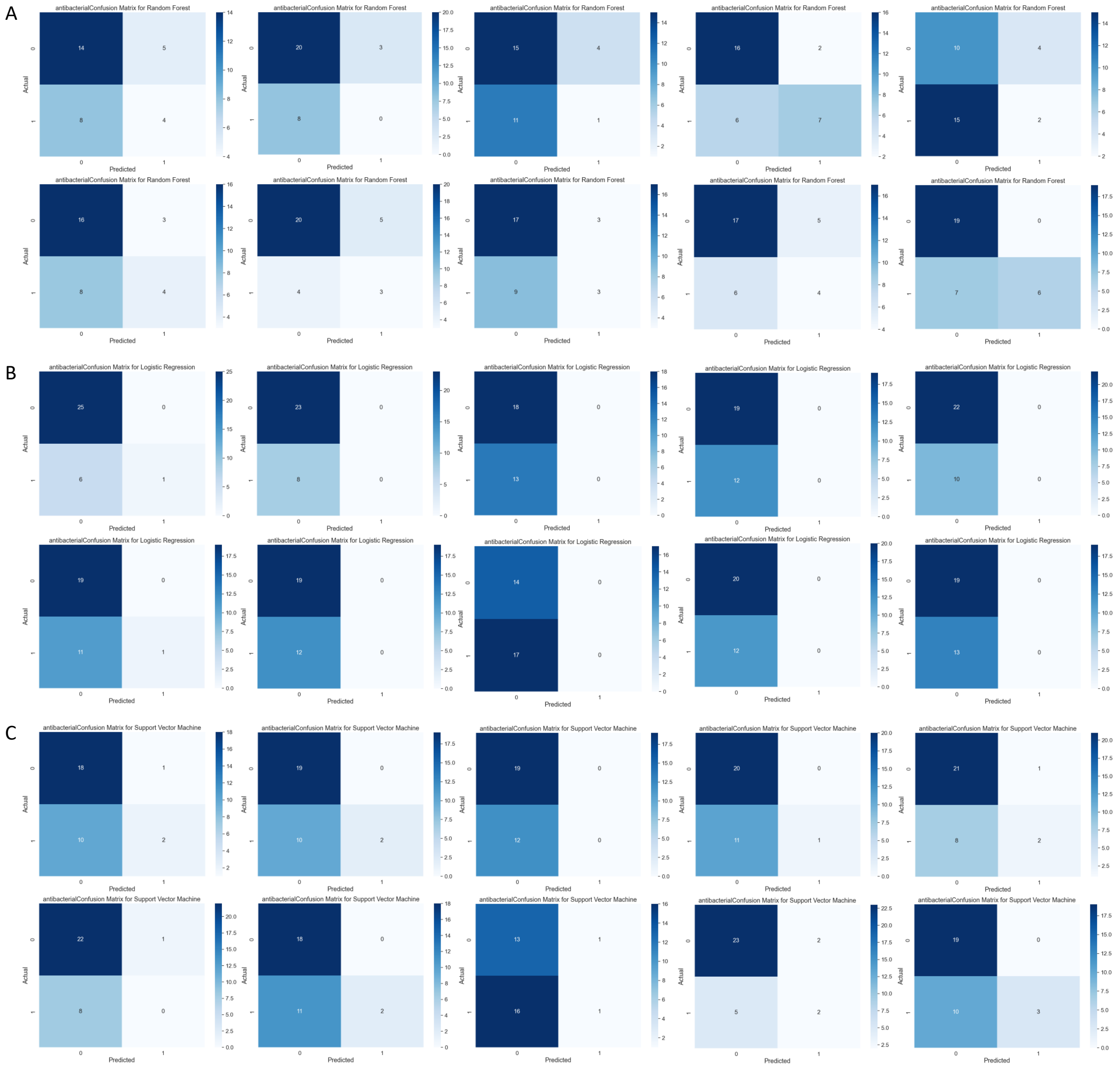


**Supplemental Figure 3**: Confusion matrices for the 10-fold cross validation for the antibacterial classification on model trained on only fungal data. A) The confusion matrices for the antibacterial random forest classifier. B) The confusion matrices for the antibacterial logistic regression classifier. C) The confusions matrices for the antibacterial support vector machine classifier.

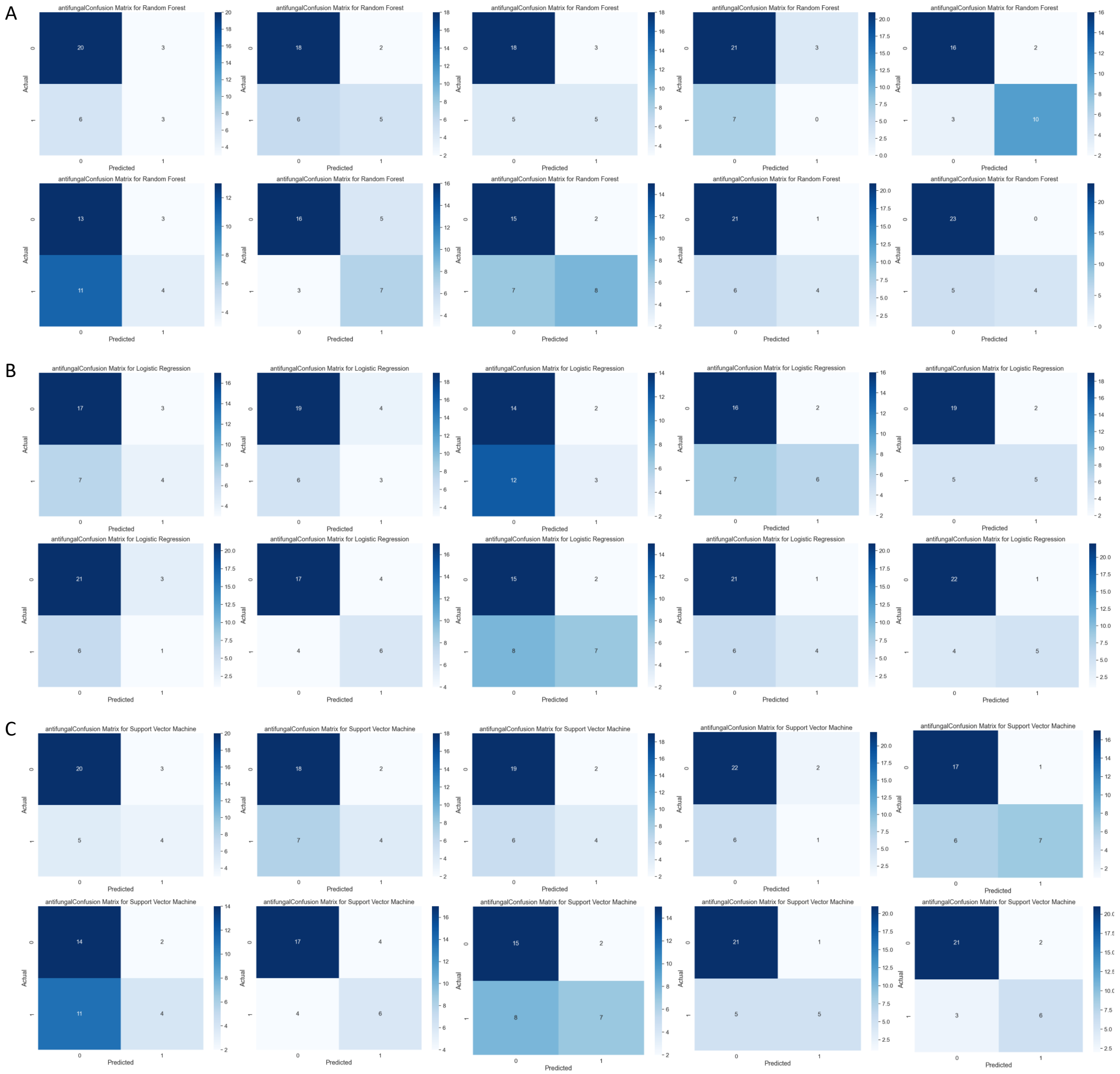


**Supplemental Figure 4**: Confusion matrices for the 10-fold cross validation for the antifungal classification on models trained on only fungal data. A) The confusion matrices for the antifungal random forest classifier. B) The confusion matrices for the antifungal logistic regression classifier. C) The confusions matrices for the antifungal support vector machine classifier.

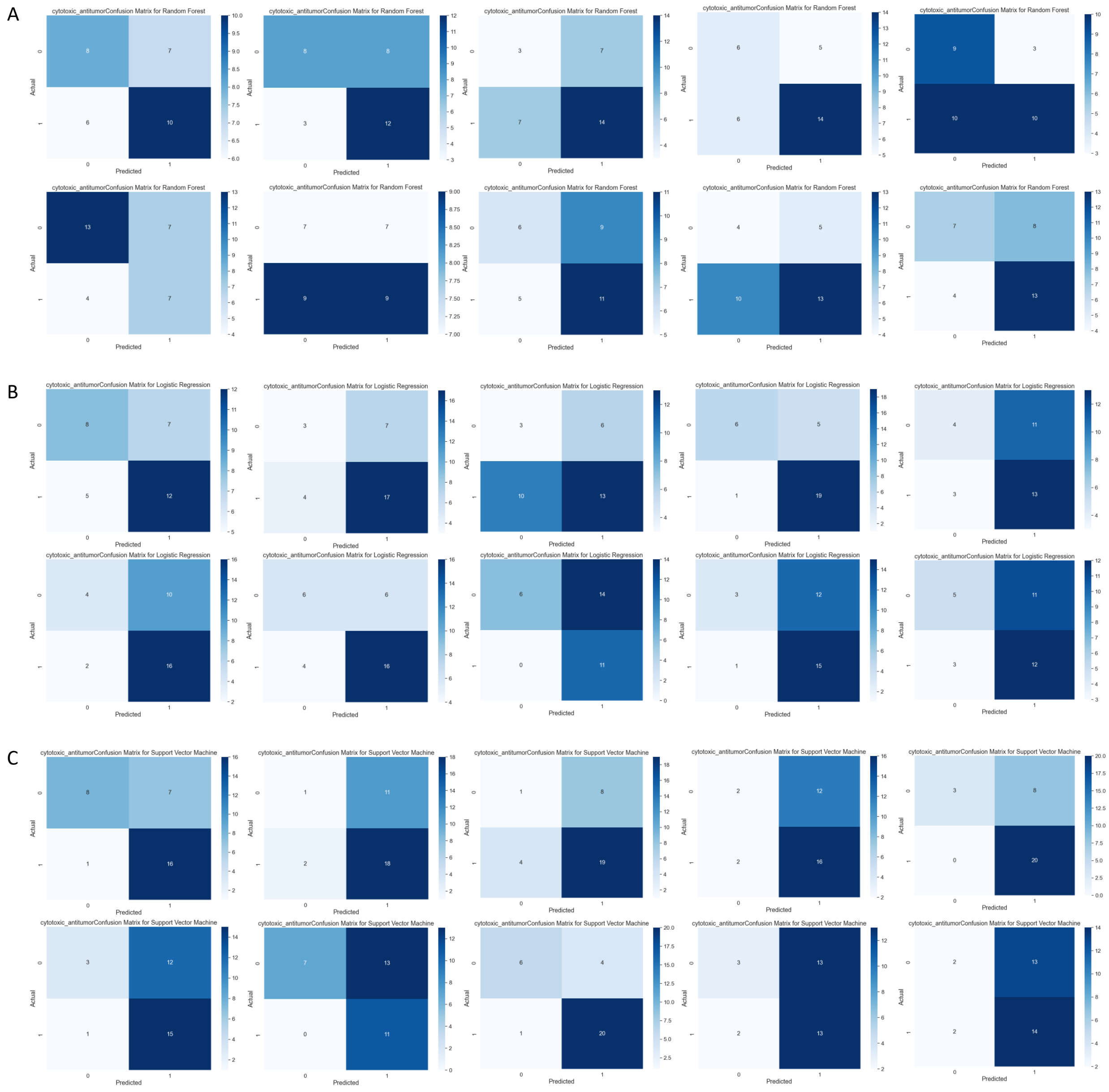


**Supplemental Figure 5**: Confusion matrices for the 10-fold cross validation for the cytotoxic/antitumor classification on models trained on only fungal data. A) The confusion matrices for the cytotoxic/antitumor random forest classifier. B) The confusion matrices for the cytotoxic/antitumor logistic regression classifier. C) The confusions matrices for the cytotoxic/antitumor support vector machine classifier.

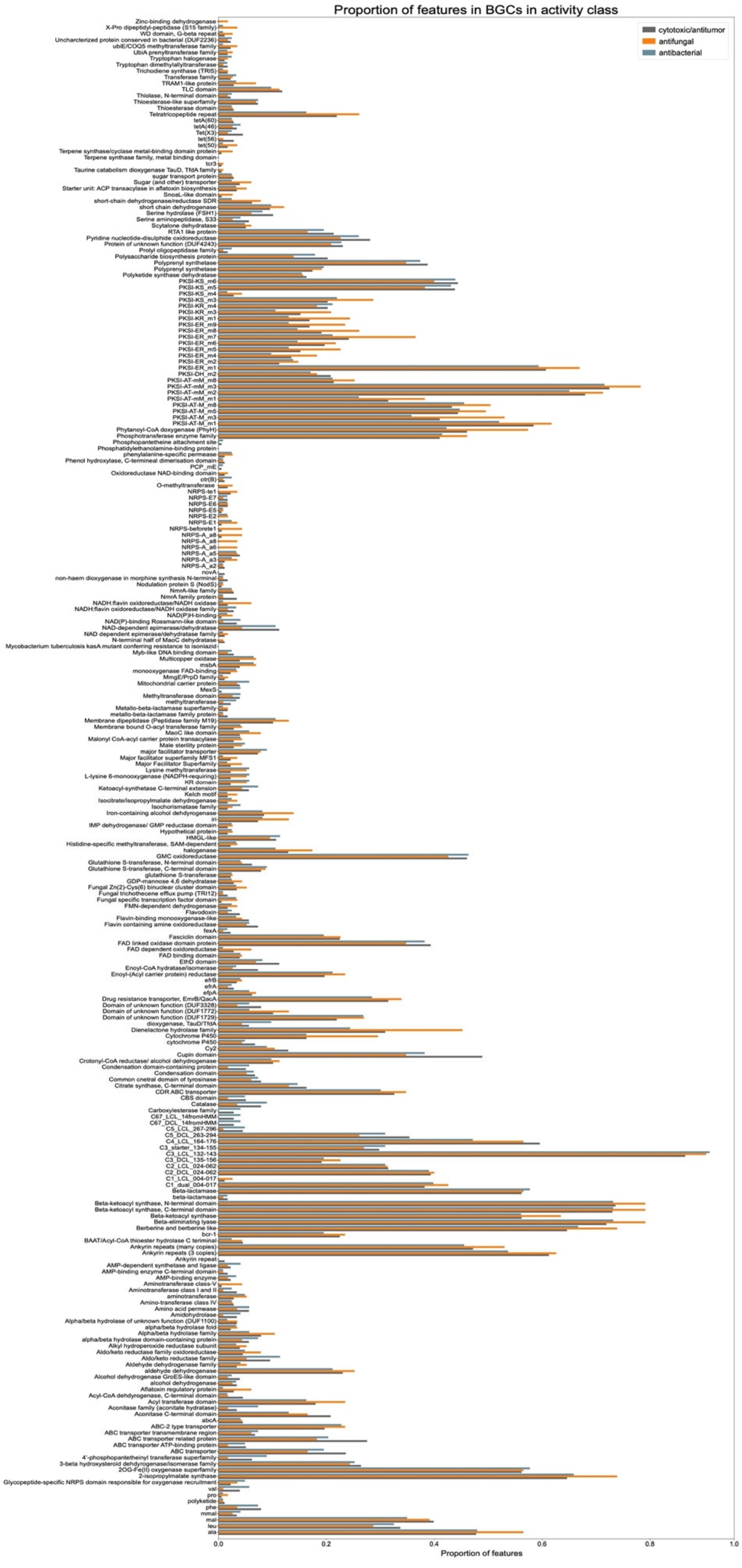

Proportion of features in BGCs in activity class
cytotoxic/antitumor
antifungal
antibacterial
Zinc-binding dehydrogenase
X-Pro dipeptidyl-peptidase (S15 family)
WD domain, G-beta repeat
Uncharacterized protein conserved in bacteria (DUF2236)
ubiE/COQ5 methyltransferase family
UbiA prenyltransferase family
Tryptophan halogenase
Tryptophan dimethylallyltransferase
Trichodiene synthase (TRI5)
Transferase family
TRAM1-like protein
TLC domain
Thiolase, N-terminal domain
Thioesterase-like superfamily
Thioesterase domain
Tetratricopeptide repeat
tetA(60)
tetA(46)
Tet(X3)
tet(56)
tet(50)
Terpene synthase/cyclase metal-binding domain protein
Terpene synthase family, metal binding domain
tcr3
Taurine catabolism dioxygenase TauD, TfdA family
sugar transport protein
Sugar (and other) transporter
Starter unit: ACP transacylase in aflatoxin biosynthesis
SnoaL-like domain
short-chain dehydrogenase/reductase SDR
short chain dehydrogenase
Serine hydrolase (FSH1)
Serine aminopeptidase, S33
Scytalone dehydratase
RTA1 like protein
Pyridine nucleotide-disulphide oxidoreductase
Protein of unknown function (DUF4243)
Prolyl oligopeptidase family
Polysaccharide biosynthesis protein
Polyprenyl synthetase
Polyprenyl synthetase
Polyketide synthase dehydratase
PKSI-KS_m6
PKSI-KS_m5
PKSI-KS_m4
PKSI-KS_m3
PKSI-KR_m4
PKSI-KR_m3
PKSI-KR_m1
PKSI-ER_m9
PKSI-ER_m8
PKSI-ER_m7
PKSI-ER_m6
PKSI-ER_m5
PKSI-ER_m4
PKSI-ER_m2
PKSI-ER_m1
PKSI-DH_m2
PKSI-AT-mM_m8
PKSI-AT-mM_m3
PKSI-AT-mM_m2
PKSI-AT-mM_m1
PKSI-AT-M_m8
PKSI-AT-M_m5
PKSI-AT-M_m3
PKSI-AT-M_m1
Phytanoyl-CoA dioxygenase (PhyH)
Phosphotransferase enzyme family
Phosphopantetheine attachment site
Phosphatidylethanolamine-binding protein
phenylalanine-specific permease
Phenol hydroxylase, C-terminal dimerisation domain
PCP_mE
Oxidoreductase NAD-binding domain
otr(B)
O-methyltransferase
NRPS-te1
NRPS-E7
NRPS-E6
NRPS-E5
NRPS-E2
NRPS-E1
NRPS-beforete1
NRPS-A_a8
NRPS-A_a8
NRPS-A_a6
NRPS-A_a5
NRPS-A_a3
NRPS-A_a2
novA
non-haem dioxygenase in morphine synthesis N-terminal
Nodulation protein S (NodS)
NmrA-like family
NmrA family protein
NADH:flavin oxidoreductase/NADH oxidase
NADH:flavin oxidoreductase/NADH oxidase family
NAD(P)H-binding
NAD(P)-binding Rossmann-like domain
NAD-dependent epimerase/dehydratase
NAD dependent epimerase/dehydratase family
N-terminal half of MaoC dehydratase
Mycobacterium tuberculosis kasA mutant conferring resistance to isoniazid
Myb-like DNA binding domain
Multicopper oxidase
msbA
monooxygenase FAD-binding
MmgE/PrpD family
Mitochondrial carrier protein
MexS
Methyltransferase domain
methyltransferase
Metallo-beta-lactamase superfamily
metallo-beta-lactamase family protein
Membrane dipeptidase (Peptidase family M19)
Membrane bound O-acyl transferase family
MaoC like domain
Malonyl CoA-acyl carrier protein transacylase
Male sterility protein
major facilitator transporter
Major facilitator superfamily MFS1
Major Facilitator Superfamily
Lysine methyltransferase
L-lysine 6-monooxygenase (NADPH-requiring)
KR domain
Ketoacyl-synthetase C-terminal extension
Kelch motif
Isocitrate/isopropylmalate dehydrogenase
Isochorismatase family
Iron-containing alcohol dehydrogenase
iri
IMP dehydrogenase/ GMP reductase domain
Hypothetical protein
HMGL-like
Histidine-specific methyltransferase, SAM-dependent
halogenase
GMC oxidoreductase
Glutathione S-transferase, N-terminal domain
Glutathione S-transferase, C-terminal domain
glutathione S-transferase
GDP-mannose 4,6 dehydratase
Fungal Zn(2)-Cys(6) binuclear cluster domain
Fungal trichothecene efflux pump (TRI12)
Fungal specific transcription factor domain
FMN-dependent dehydrogenase
Flavodoxin
Flavin-binding monooxygenase-like
Flavin containing amine oxidoreductase
fexA
Fasciclin domain
FAD linked oxidase domain protein
FAD dependent oxidoreductase
FAD binding domain
EthD domain
Enoyl-CoA hydratase/isomerase
Enoyl-(Acyl carrier protein) reductase
efrB
efrA
efpA
Drug resistance transporter, EmrB/QacA
Domain of unknown function (DUF3328)
Domain of unknown function (DUF1772)
Domain of unknown function (DUF1729)
dioxygenase, TauD/TfdA
Dienelactone hydrolase family
Cytochrome P450
cytochrome P450
Cy2
Cupin domain
Crotonyl-CoA reductase/ alcohol dehydrogenase
Condensation domain-containing protein
Condensation domain
Common central domain of tyrosinase
Citrate synthase, C-terminal domain
CDR ABC transporter
CBS domain
Catalase
Carboxylesterase family
C67_LCL_14fromHMM
C67_DCL_14fromHMM
C5_LCL_267-296
C5_DCL_263-294
C4_LCL_164-176
C3_starter_134-155
C3_LCL_132-143
C3_DCL_135-156
C2_LCL_024-062
C2_DCL_024-062
C1_LCL_004-017
C1_dual_004-017
Beta-lactamase
beta-lactamase
Beta-ketoacyl synthase, N-terminal domain
Beta-ketoacyl synthase, C-terminal domain
Beta-ketoacyl synthase
Beta-eliminating lyase
Berberine and berberine like
bcr-1
BAAT/Acyl-CoA thioester hydrolase C terminal
Ankyrin repeats (many copies)
Ankyrin repeats (3 copies)
Ankyrin repeat
AMP-dependent synthetase and ligase
AMP-binding enzyme C-terminal domain
AMP-binding enzyme
Aminotransferase class-V
Aminotransferase class I and II
aminotransferase
Amino-transferase class IV
Amino acid permease
Amidohydrolase
Alpha/beta hydrolase of unknown function (DUF1100)
alpha/beta hydrolase fold
Alpha/beta hydrolase family
alpha/beta hydrolase domain-containing protein
Alkyl hydroperoxide reductase subunit
Aldo/keto reductase family oxidoreductase
Aldo/keto reductase family
Aldehyde dehydrogenase family
aldehyde dehydrogenase
Alcohol dehydrogenase GroES-like domain
alcohol dehydrogenase
Aflatoxin regulatory protein
Acyl-CoA dehydrogenase, C-terminal domain
Acyl transferase domain
Aconitase family (aconitate hydratase)
Aconitase C-terminal domain
abcA
ABC-2 type transporter
ABC transporter transmembrane region
ABC transporter related protein
ABC transporter ATP-binding protein
ABC transporter
4'-phosphopantetheinyl transferase superfamily
3-beta hydroxysteroid dehydrogenase/isomerase family
2OG-Fe(II) oxygenase superfamily
2-isopropylmalate synthase
Glycopeptide-specific NRPS domain responsible for oxygenase recruitment
val
pro
polyketide
phe
mmal
mal
leu
ala
0.0
0.2
0.4
0.6
0.8
1.0
Proportion of features

**Supplemental Figure 6**: Proportion of all features in BGCs in each activity class for the fungal dataset. X-axis displays the proportion of features in each activity class: cytotoxic/antitumor (grey), antifungal (orange), antibacterial (blue) and Y-axis shows the features.

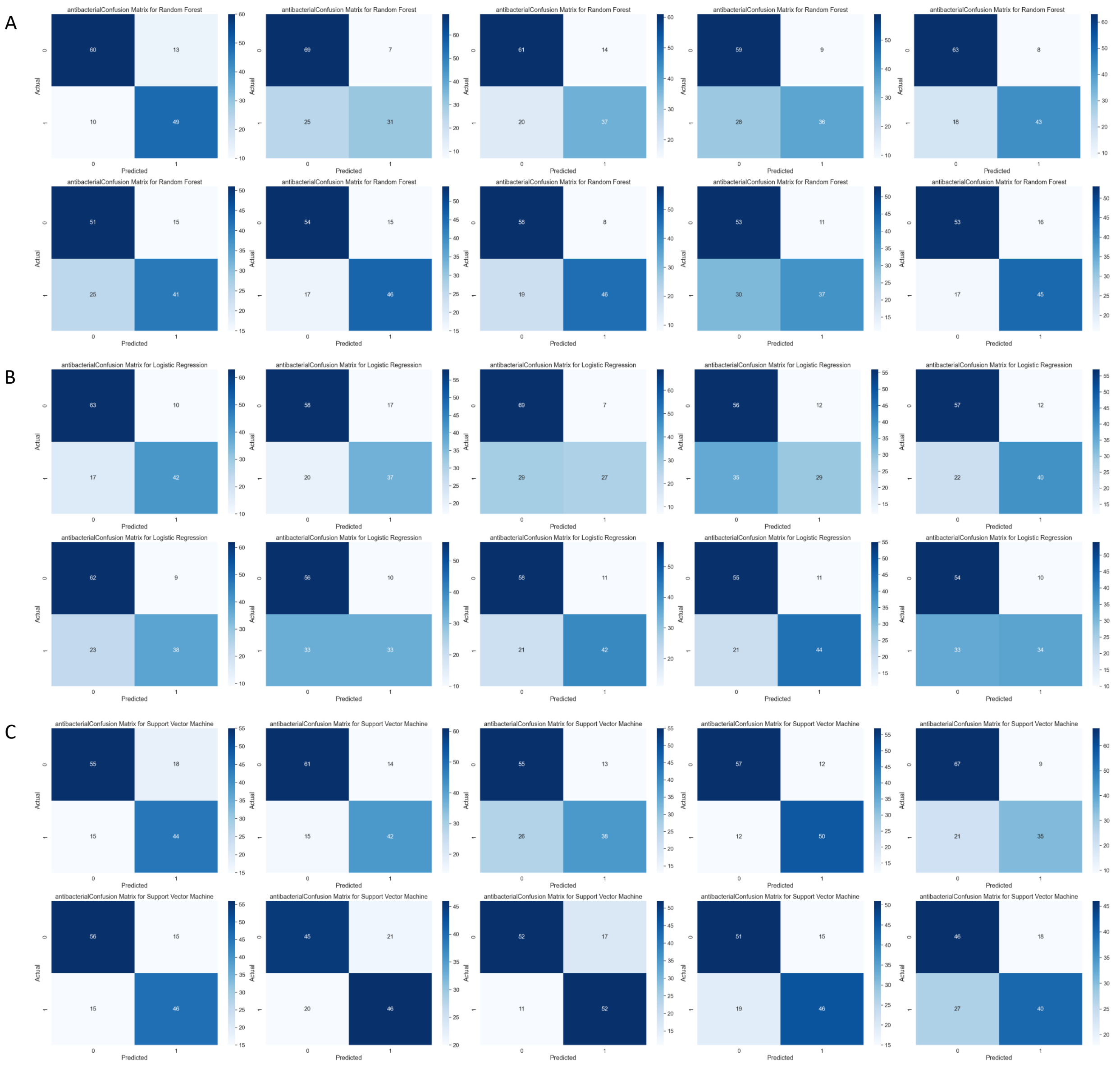


**Supplemental Figure 7**: Confusion matrices for the 10-fold cross validation for the antibacterial classification on models trained on both fungal and bacterial data. A) The confusion matrices for the antibacterial random forest classifier. B) The confusion matrices for the antibacterial logistic regression classifier. C) The confusions matrices for the antibacterial support vector machine classifier.

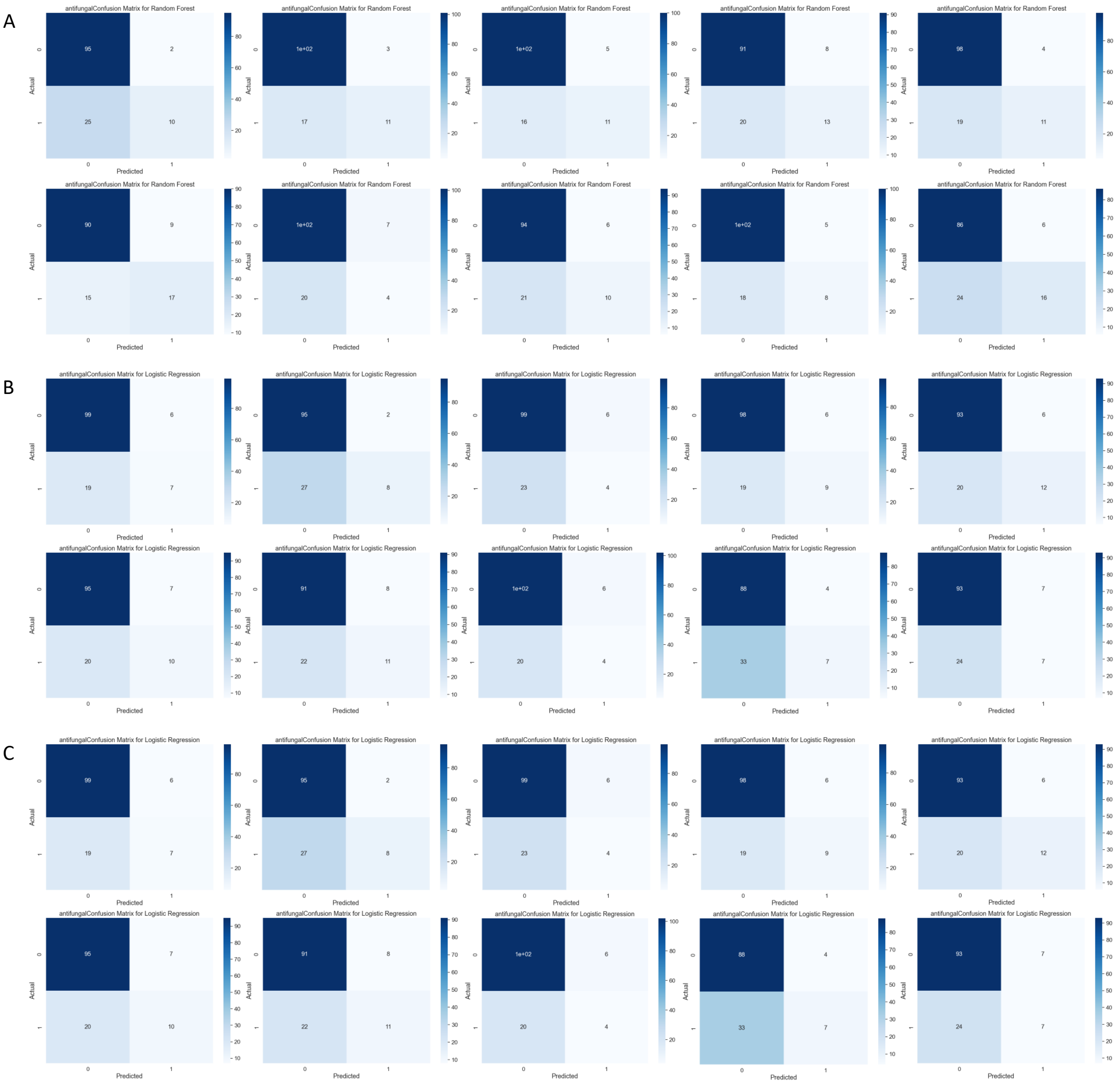


**Supplemental Figure 8**: Confusion matrices for the 10-fold cross validation for the antifungal classification on models trained on both fungal and bacterial data. A) The confusion matrices for the antifungal random forest classifier. B) The confusion matrices for the antifungal logistic regression classifier. C) The confusions matrices for the antifungal support vector machine classifier.

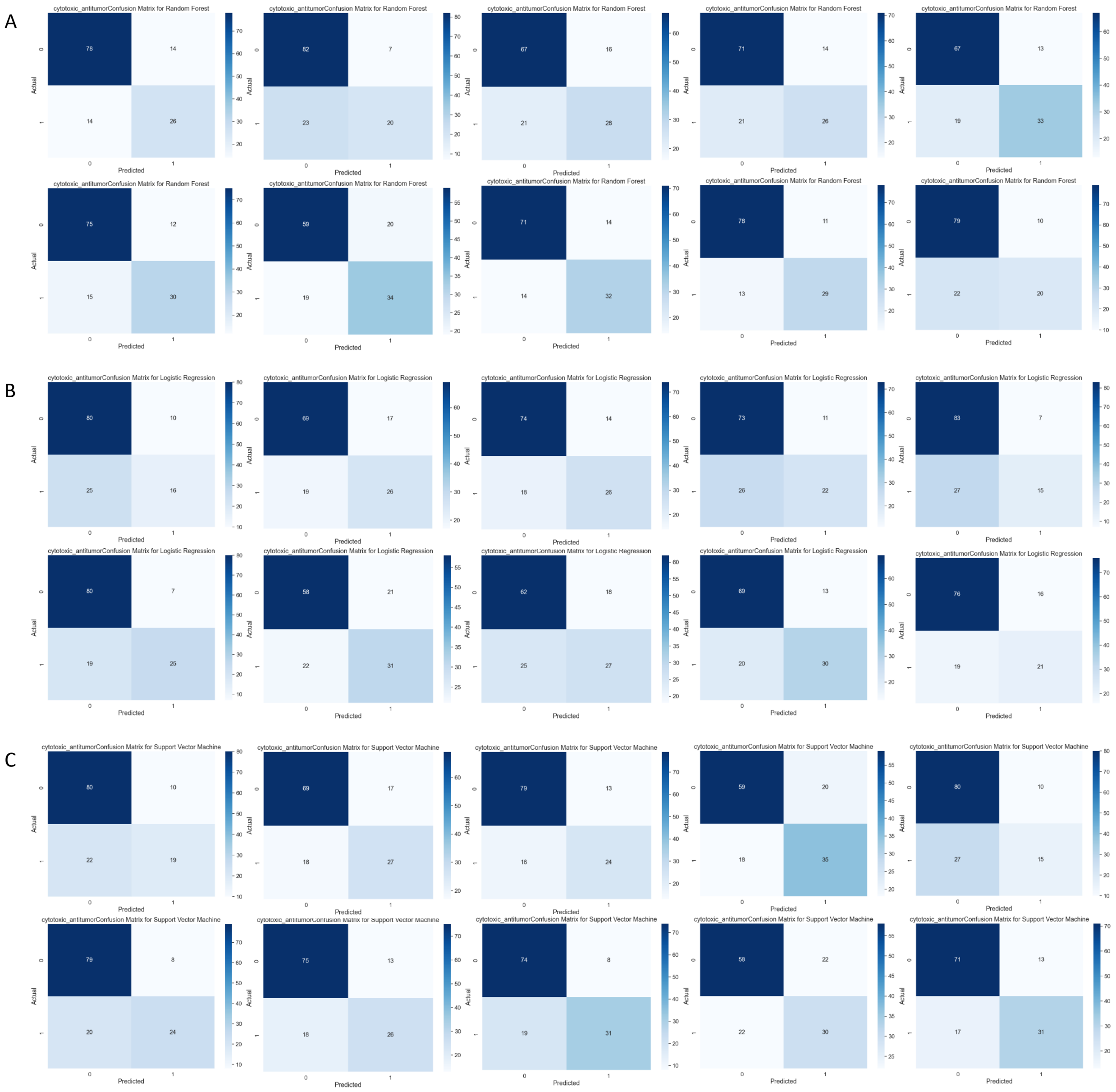


**Supplemental Figure 9**: Confusion matrices for the 10-fold cross validation for the cytotoxic/antitumor classification on models trained on both fungal and bacterial data. A) The confusion matrices for the cytotoxic/antitumor random forest classifier. B) The confusion matrices for the cytotoxic/antitumor logistic regression classifier. C) The confusions matrices for the cytotoxic/antitumor support vector machine classifier.

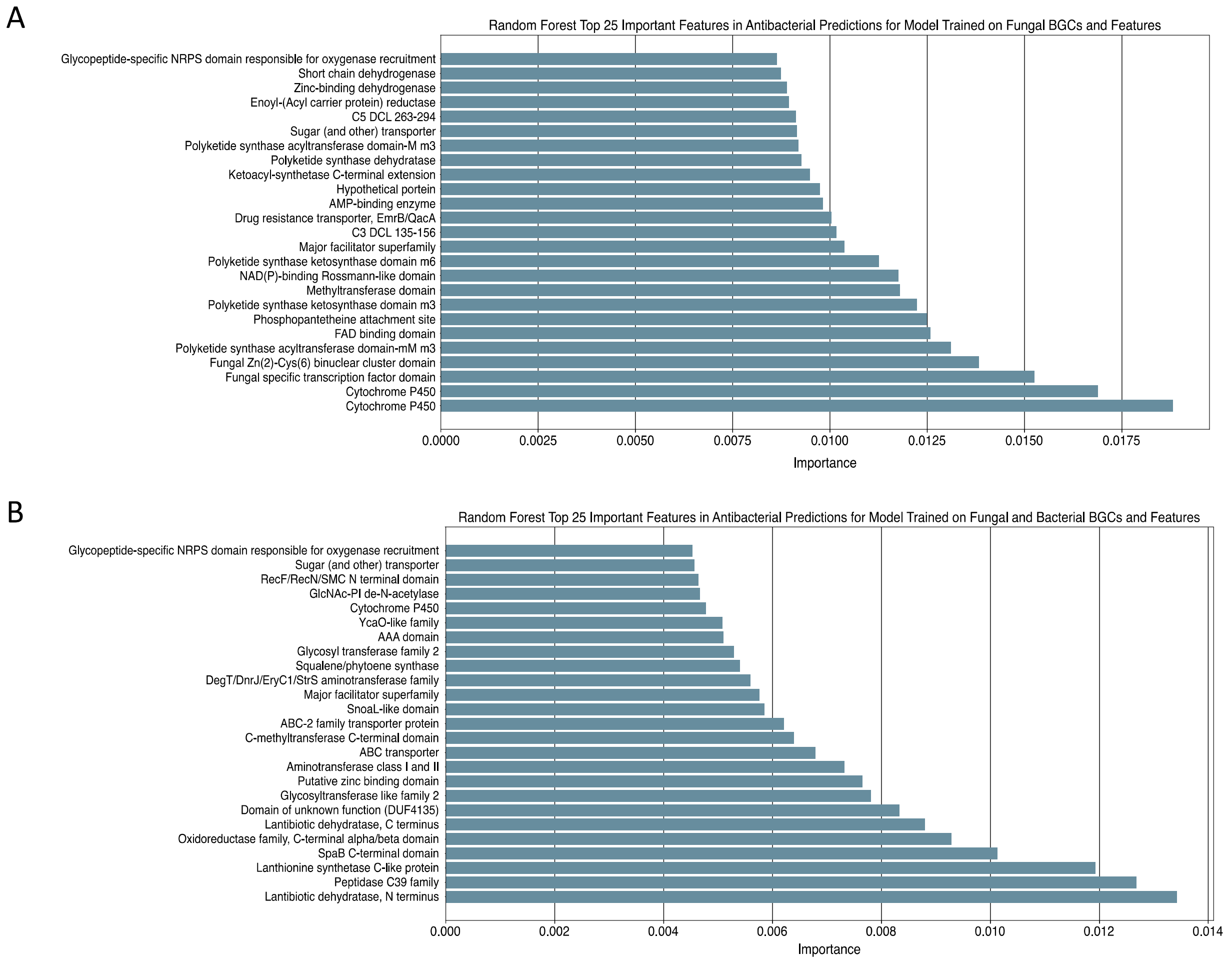


**Supplemental Figure 10**: A) Top 25 important features in antibacterial predictions for the random forest trained on fungal data. B) Top 25 important features in antibacterial predictions for the random forest model trained on fungal and bacterial data. X-axis shows importance and Y-axis shows the features.

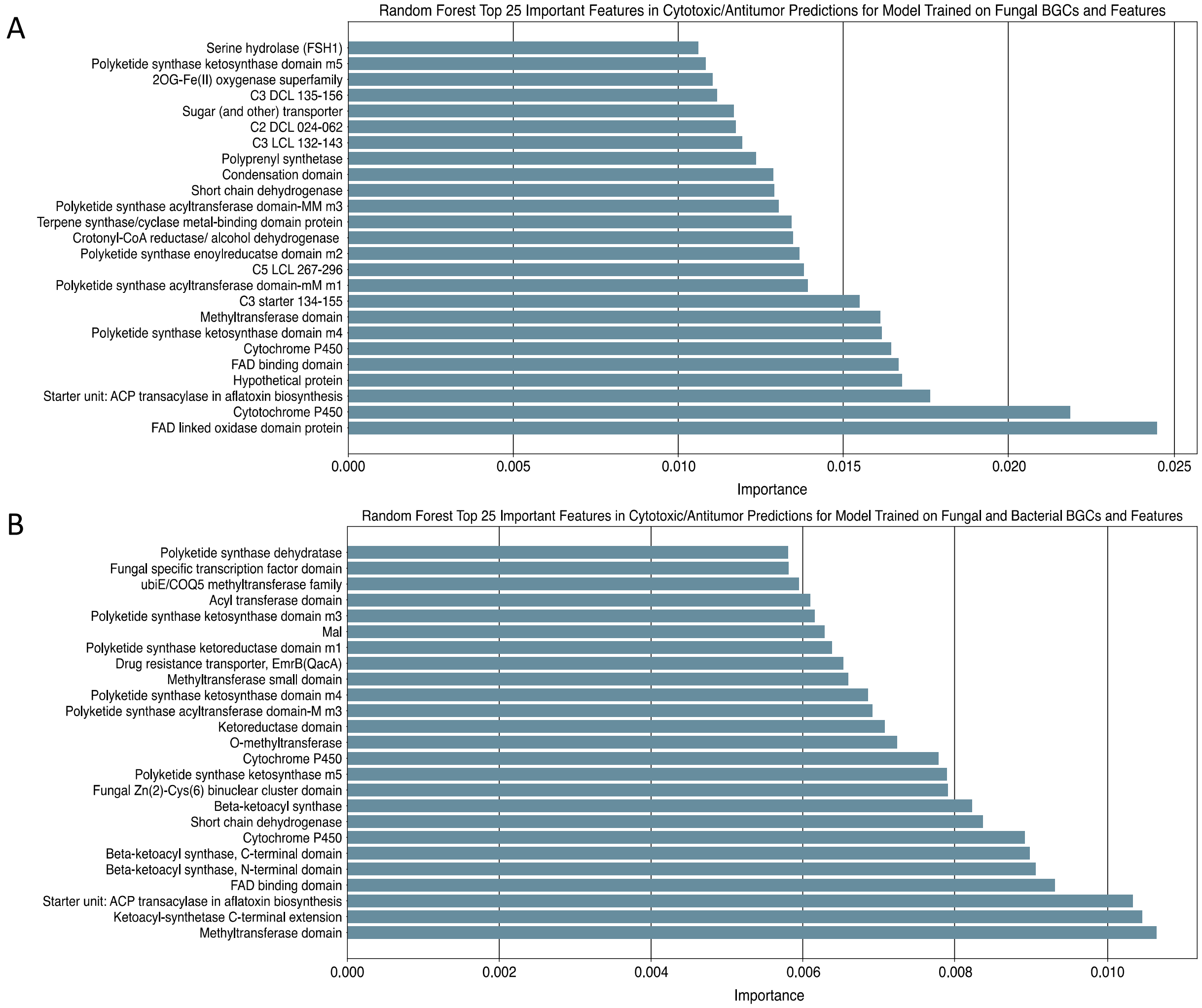


**Supplemental Figure 11**: A) Top 25 important features in cytotoxic/antitumor predictions for the random forest model trained on fungal data. B) Top 25 important features in cytotoxic/antitumor predictions for the random forest model trained on fungal and bacterial data. X-axis shows importance and Y-axis shows the features.

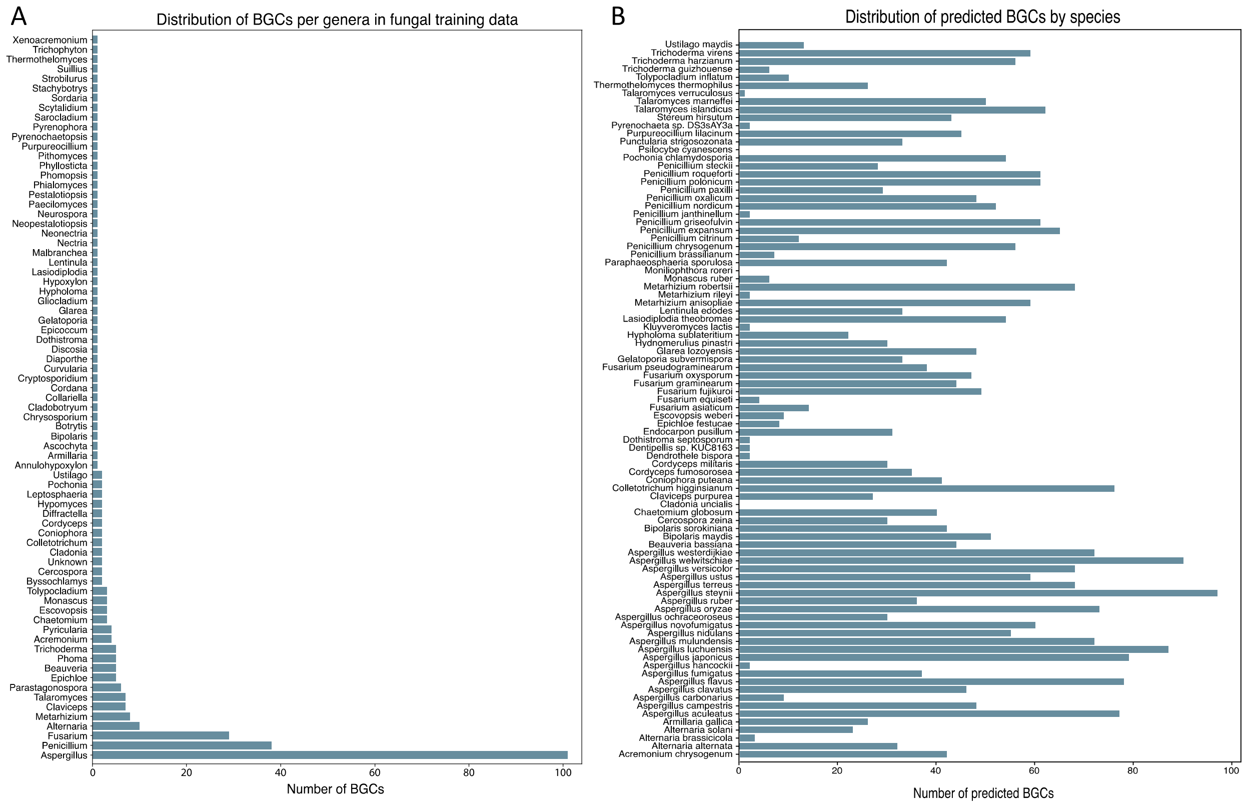


**Supplemental Figure 12**: A) Distribution of biosynthetic gene clusters (BGCs) in training dataset. X axis shows the number of BGCs, and Y-axis shows the genera in the training dataset. B) Distribution of predicted BGCs by species in the training dataset.

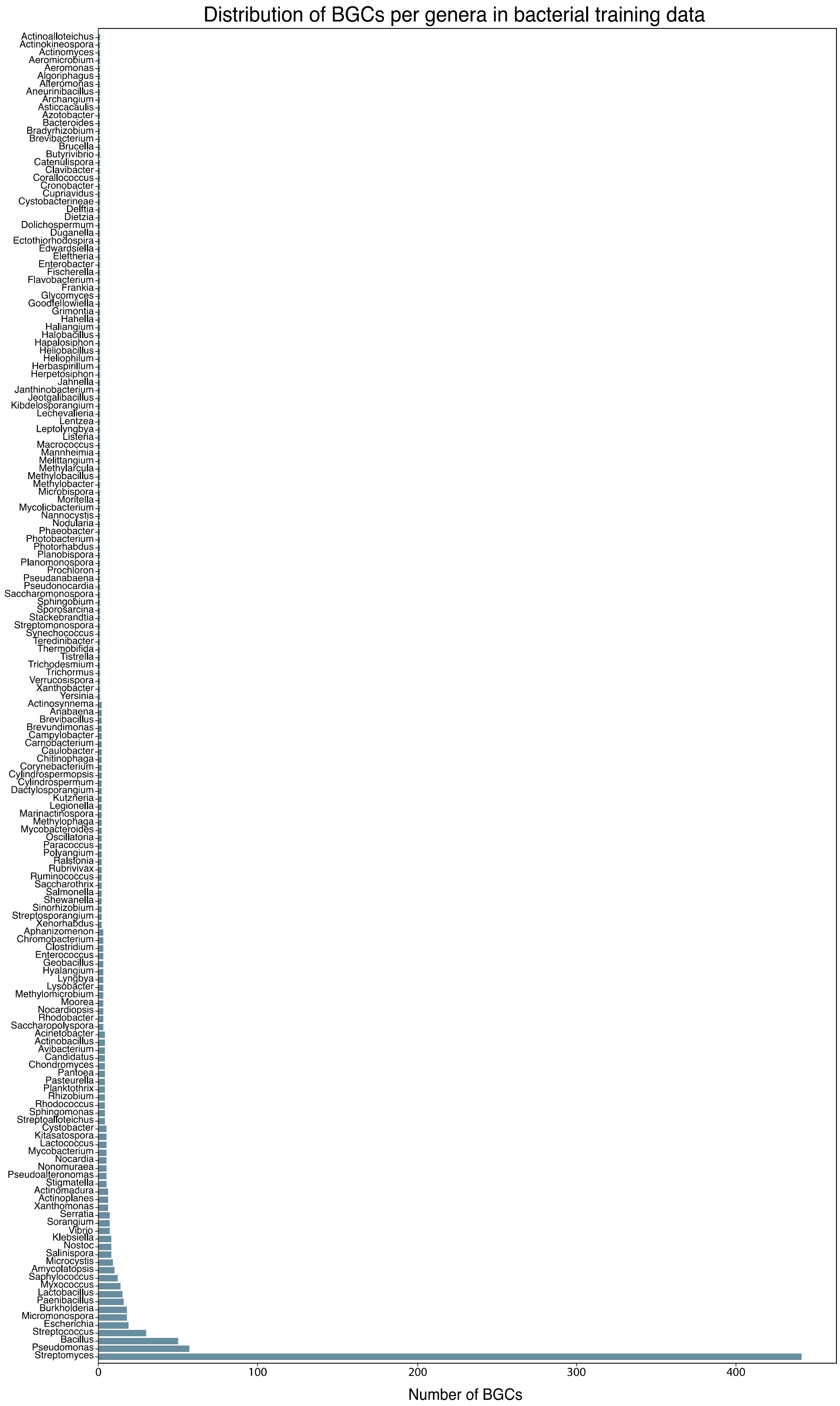
Distribution of BGCs per genera in bacterial training data
Actinoalloteichus
Actinokineospora
Actinomyces
Aeromicrobium
Aeromonas
Algoriphagus
Alteromonas
Aneurinibacillus
Archangium
Asticcacaulis
Azotobacter
Bacteroides
Bradyrhizobium
Brevibacterium
Brucella
Butyrivibrio
Catenulispora
Clavibacter
Corallococcus
Cronobacter
Cupriavidus
Cystobacterineae
Delftia
Dietzia
Dolichospermum
Duganella
Ectothiorhodospira
Edwardsiella
Eleftheria
Enterobacter
Fischerella
Flavobacterium
Frankia
Glycomyces
Goodfellowiella
Grimontia
Hahella
Haliangium
Halobacillus
Hapalosiphon
Heliobacillus
Heliophilum
Herbaspirillum
Herpetosiphon
Jahnella
Janthinobacterium
Jeotgalibacillus
Kibdelosporangium
Lechevalieria
Lentzea
Leptolyngbya
Listeria
Macrococcus
Mannheimia
Melittangium
Methylarcula
Methylobacillus
Methylobacter
Microbispora
Moritella
Mycolicibacterium
Nannocystis
Nodularia
Phaeobacter
Photobacterium
Photorhabdus
Planobispora
Planomonospora
Prochloron
Pseudanabaena
Pseudonocardia
Saccharomonospora
Sphingobium
Sporosarcina
Stackebrandtia
Streptomonospora
Synechococcus
Teredinibacter
Thermobifida
Tistrella
Trichodesmium
Trichormus
Verrucosispora
Xanthobacter
Yersinia
Actinosynnema
Anabaena
Brevibacillus
Brevundimonas
Campylobacter
Carnobacterium
Caulobacter
Chitinophaga
Corynebacterium
Cylindrospermopsis
Cylindrospermum
Dactylosporangium
Kutzneria
Legionella
Marinactinospora
Methylophaga
Mycobacteroides
Oscillatoria
Paracoccus
Polyangium
Ralstonia
Rubrivivax
Ruminococcus
Saccharothrix
Salmonella
Shewanella
Sinorhizobium
Streptosporangium
Xenorhabdus
Aphanizomenon
Chromobacterium
Clostridium
Enterococcus
Geobacillus
Hyalangium
Lyngbya
Lysobacter
Methylomicrobium
Moorea
Nocardiopsis
Rhodobacter
Saccharopolyspora
Acinetobacter
Actinobacillus
Avibacterium
Candidatus
Chondromyces
Pantoea
Pasteurella
Planktothrix
Rhizobium
Rhodococcus
Sphingomonas
Streptoalloteichus
Cystobacter
Kitasatospora
Lactococcus
Mycobacterium
Nocardia
Nonomuraea
Pseudoalteromonas
Stigmatella
Actinomadura
Actinoplanes
Xanthomonas
Serratia
Sorangium
Vibrio
Klebsiella
Nostoc
Salinispora
Microcystis
Amycolatopsis
Saphylococcus
Myxococcus
Lactobacillus
Paenibacillus
Burkholderia
Micromonospora
Escherichia
Streptococcus
Bacillus
Pseudomonas
Streptomyces
0
100
200
300
400
Number of BGCs

**Supplemental Figure 13**: Distribution of biosynthetic gene clusters (BGCs) in bacterial training dataset. X axis shows the number of BGCs and Y-axis shows the genera in the training dataset.

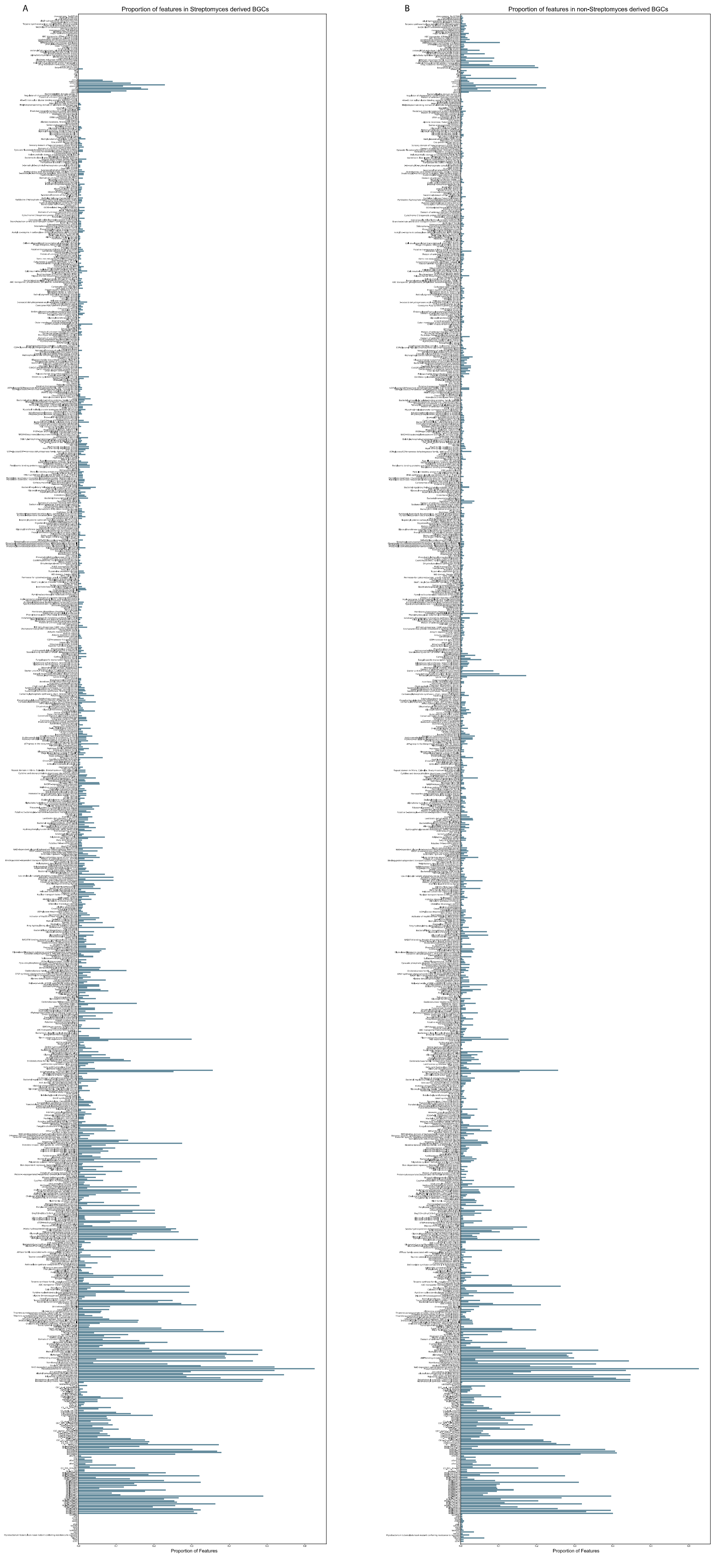
A
Proportion of features in Streptomyces derived BGCs
Proportion of Features
B
Proportion of features in non-Streptomyces derived BGCs
Proportion of Features

**Supplemental Figure 14**: Proportion of all features in the fungal and bacterial dataset. A) Proportion of features present in *Streptomyces* the derived BGCs. B) Proportion of features present in non-*Streptomyces* derived BGCs.
